# Pigment-dispersing factor neuropeptides act as multifunctional hormones and modulators in tardigrades

**DOI:** 10.1101/2024.07.30.605422

**Authors:** Soumi Dutta, Lars Hering, Milena M. Grollmann, Niklas Metzendorf, Vladimir Gross, Kazuharu Arakawa, Susanne Neupert, Monika Stengl, Friedrich W. Herberg, Georg Mayer

**Affiliations:** Department of Zoology, Institute of Biology, University of Kassel, Germany; BMBF Research Initiative for the Conservation of Biodiversity (FEdA), Senckenberg Nature Research Society, Frankfurt am Main, Germany; Institute for Advanced Biosciences, Keio University, Tsuruoka City, Yamagata, Japan; Department of Animal Physiology/Neuroethology, Institute of Biology, University of Kassel, Germany; Department of Biochemistry, Institute of Biology, University of Kassel, Germany

## Abstract

Pigment-dispersing factors (PDFs) are neuropeptides that play key roles in controlling the circadian rhythms in various insects, whereas their function remains elusive in other protostomes including tardigrades (water bears). Here we show that the three PDFs of the tardigrade *Hypsibius exemplaris* are co-localized in two pairs of inner lobe cells in the brain, whereas only one PDF occurs in four additional cerebral and two extracerebral cells. The axons of the inner lobe cells pass through the contralateral brain hemisphere, descend to the ventral nerve cord and terminate in two pairs of potential release sites in the posteriormost trunk ganglion. Using *in vitro* assays, we demonstrate that all three PDFs and their deorphanized receptor (PDFR) are functional. Widespread localization of PDFR suggests that tardigrade PDFs may act as multifunctional hormones and neuromodulators that control major functions including light detection, neural processing, locomotion, feeding, digestion, osmoregulation, growth, embryonic development, and oogenesis/reproduction.

## Introduction

Pigment-dispersing factors (PDFs) are conserved neuropeptides that were initially identified in crustaceans as hormones (PDHs) that control epidermal color changes and visual adaptation to light^1–3^. Subsequent studies suggested that PDFs play key roles in regulating the rest-activity rhythms as circadian clock outputs in that they directly couple bilateral clock centers in the two brain hemispheres^4–7^. They additionally control infradian rhythms of photoperiodic responses in different insects^8,9^. The circadian functions of PDFs in insects resemble those of the vasoactive intestinal peptide (VIP) in vertebrates^10^, although the *pdf* and *vip* lineages are no sister groups and the ancestral *pdf* gene most likely evolved in protostomes^11^. While the last common ancestor of protostomes possessed only one *pdf* gene, a duplication might have led to two homologs in the last common ancestor of ecdysozoans (molting animals), followed by a subsequent loss of one of them in tardigrades (water bears) and arthropods (Fig. 1a). Repeated duplications likely gave rise to multiple *pdf* copies in tardigrades and decapod crustaceans^12^.

**Fig. 1.**
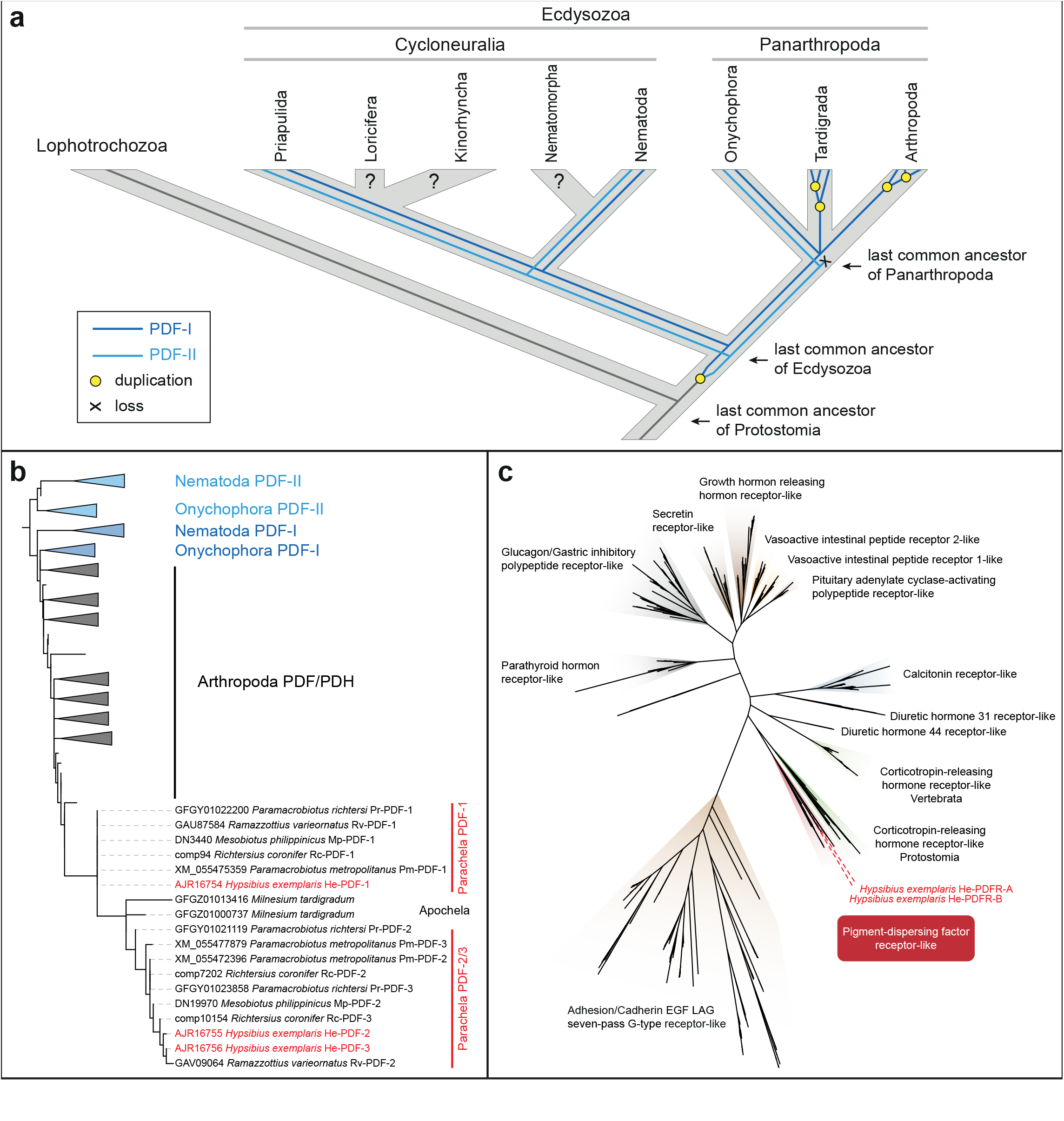
**Evolutionary history of PDF/PDH peptides in ecdysozoans and phylogeny of PDF peptides and PDF receptor proteins**. (**a**) Scenario on evolution of PDF peptides in ecdysozoans. Note presence of two PDF peptides in last common ancestors of Panarthropoda and Ecdysozoa. Modified from Mayer et al.^12^. (**b**) Maximum likelihood tree of PDFs across ecdysozoans. Note position of monophyletic eutardigrade PDFs nested within arthropod PDF/PDH clade and occurrence of two distinct clades of parachelan PDFs (PDF-1 and PDF- 2/3). PDF peptides of *H. exemplaris* are highlighted in red. (**c**) Maximum likelihood tree of ∼1,000 class B G protein-coupled receptor proteins (GPCRs). Dataset was obtained previously^11^ from cluster analysis of ∼18,000 bilaterian GPCRs. Both PDFR splice variants (He-PDFR-A and He-PDFR-B) of *H. exemplaris* fall into monophyletic group of PDF receptors of protostomes (highlighted in red).

Most information about the localization of PDFs is available from insects^4,5,13,14^ but there are also some data from crustaceans^15–18^, onychophorans (velvet worms)^11,12^, nematodes^19^ and mollusks^20^. The PDF-immunoreactive (PDF*-ir*) cells of insects are arranged in small clusters associated with the optic lobe neuropils of the compound eyes^4–6,9,21,22^ and additional somata occur in the protocerebrum of some species studied to date^8,23,24^. PDF*-ir* neurons associated with the visual system arborize in a small neuropil, the accessory medulla, which was identified as a pacemaker center controlling circadian locomotor activity rhythms^25^. A similar arrangement has been reported from crustaceans, but the existence of accessory medullae in this group and their involvement in the circadian clock have not been demonstrated^16,17,26^.

While data on PDF immunoreactivity are unavailable from other arthropods, such as chelicerates (spiders and allies) and myriapods (e.g., centipedes and millipedes), there is no evidence of accessory medullae in onychophorans^11,12,27^, which along with tardigrades represent the closest living relatives of arthropods (Fig. 1a). Hence, there are no PDF*-ir* somata directly associated with the onychophoran eyes that are most likely homologous with the median ocelli rather than the compound eyes of arthropods^27^. Despite the lack of accompanying PDF-*ir* somata, the visual neuropil of each onychophoran eye does contain numerous PDF-I*-ir* and PDF-II*-ir* fibers^11^, most likely originating from a specific subset of protocerebral neurons. Due to their axonal contiguity with the photoreceptors, which exhibit maximum sensitivity to blue light (∼480 nm)^28^, these neurons are the most likely candidates for circadian pacemakers. Besides in the protocerebrum, numerous PDF-*ir* somata occur throughout the remaining central nervous system of onychophorans, including the deutocerebrum, the circumpharyngeal nerve cords, and the ventral nerve cords^11,12^. All PDF- *ir* neurons of onychophorans invariably contain both peptides, PDF-I and PDF-II, though these seem to be expressed at different levels in different groups of neurons^11^.

Like onychophorans, the nematode *Caenorhabditis elegans* exhibits both ancestral *pdf* genes^12^, *pdf-I* and *pdf-II* (Fig 1a), whose products are localized in different neurons including sensory, motor, mechanosensory and interneurons^19^. Beyond this, both peptides occur in tissues and cells outside the nervous system, including those of rectal glands, the intestino- rectal valve, and the arcade cells associated with the pharynx^19^. However, at least in the nervous system of *Caenorhabditis elegans*, PDF-I and PDF-II are not entirely co-localized^19^. Interestingly, the PDF receptor (PDFR) of nematodes is expressed in several tissues, suggesting that PDF-I and PDF-II may act as hormones that control multiple functions in these animals, including locomotion, chemosensation, mechanosensation, and integration of other external stimuli^19,29–31^. Similar observations have been made in onychophorans, in which at least some PDFR-*ir* and PDF-*ir* cells are spatially separated^11^ and there are potential release sites into the lumen of the heart^12^, suggesting hormonal release into the hemolymph and a dual role of PDFs as hormones and neuromodulators in velvet worms.

Despite their key phylogenetic position as members of Panarthropoda and Ecdysozoa (Fig. 1a), virtually nothing is known about the organization of the PDF/PDFR system in tardigrades. Three *pdf* homologs have been identified previously^12,32^ in the genome and transcriptomes of the model eutardigrade^33,34^ *Hypsibius exemplaris*, but their localization and functionalities are unknown. A cross-reactive antiserum, raised against the synthetic β-PDH peptide of the crustacean *Uca pugilator*^15^ and commonly used for localizing PDF*-ir* neurons in various insects^4,5,13,21^, crustaceans^15–17^, onychophorans^12^, and even mollusks^20^, previously failed to detect the PDFs of tardigrades^12^. One *pdfr* ortholog was further identified *in silico* in the genome assemblies and predicted proteomes of two eutardigrade species, including *H. exemplaris*^32^, but the existence of potential isoforms, their functionality and localization remain unexplored.

The present study thus sets out to close these gaps and to address the following questions: (i) Are the three PDFs of *H. exemplaris* functional? (ii) Where are they localized and are any of them co-localized? (iii) How does the position of PDF-*ir* neurons and their projections relate to that in other protostomes, in particular insects and crustaceans? (iv) Are there differences in expression levels between the three *pdf* genes? (v) How do these compare to the expression levels of *pdfr* and its potential isoforms? (vi) Is PDFR co-localized with PDFs? If yes, this would suggest autoreception, like in insects^35–38^, whereas if PDFR is localized in additional cells, this would indicate hormonal release of PDFs into the hemolymph, like in crustaceans, onychophorans, and nematodes^11,18,19^. Clarifying these questions will provide insights into the evolutionary changes in the organization and function of PDF/PDFR signaling across panarthropods and ecdysozoans.

## Results

### Orthology, composition and genomic location of *pdf* genes in eutardigrades

Genomic searches using three previously identified *pdf* sequences from the transcriptomes of *H. exemplaris*^12^ as queries confirmed the existence of three *pdf* genes in this tardigrade species: *He-pdf-1*, *He-pdf-2*, and *He-pdf-3*. To clarify orthology of identified homologs, we performed a phylogenetic analysis of *pdf* genes across eutardigrades. Genomic and transcriptomic screening, including reciprocal BLAST searches, revealed two to three *pdf* homologs in several species of Parachela (two in *Ramazzottius varieornatus* and *Mesobiotus philippinicus*, and three in *Richtersius coronifer*, *Paramacrobiotus richtersi* and *Paramacrobiotus metropolitanus*), whereas *Milnesium tardigradum* from the Apochela clade has two *pdf* homologs (Fig. 1b; Supplementary Fig. 1).

We further analyzed the structure and arrangement of *pdf* genes in the assembled genomes of *H. exemplaris* and *Ramazzottius varieornatus*. The analysis revealed that *He*-*pdf*-*1* of *H. exemplaris* (574 bp) is located on the forward strand of scaffold0024 (MTYJ01000024.1), whereas *He*-*pdf*-*2* (1,401 bp) and *He*-*pdf*-*3* (643 bp) are located on the forward and reverse strands of scaffold0058 (MTYJ01000058.1), respectively (Fig. 2a; Supplementary Data 1). *He*-*pdf*-*1* consists of three exons (111 bp, 109 bp and 35 bp) separated by two introns (126 bp and 193 bp), while each of the two other *pdf* genes, *He*-*pdf*- *2* and *He*-*pdf*-*3*, comprise only two exons (*He*-*pdf*-*2*: 117 bp and 177 bp; *He*-*pdf*-*3*: 117 bp and 222 bp) separated by one intron (*He*-*pdf*-*2*: 1107 bp; *He*-*pdf*-*3*: 304 bp). In *Ramazzottius varieornatus*, the *Rv-pdf*-*1* gene (469 bp) is located on the forward strand of scaffold001 (BDGG01000001.1) and *Rv-pdf*-*2* (440 bp) on the reverse strand of scaffold020 (BDGG01000020.1), respectively. Each gene consists of two exons (*Rv*-*pdf*-*1*: 111 bp and 147 bp; *Rv*-*pdf*-*2*: 117 bp and 225 bp) separated by one intron (*Rv*-*pdf*-*1*: 211 bp; *Rv*-*pdf*-*2*: 98 bp) (Fig. 2a; Supplementary Data 1).

**Fig. 2.**
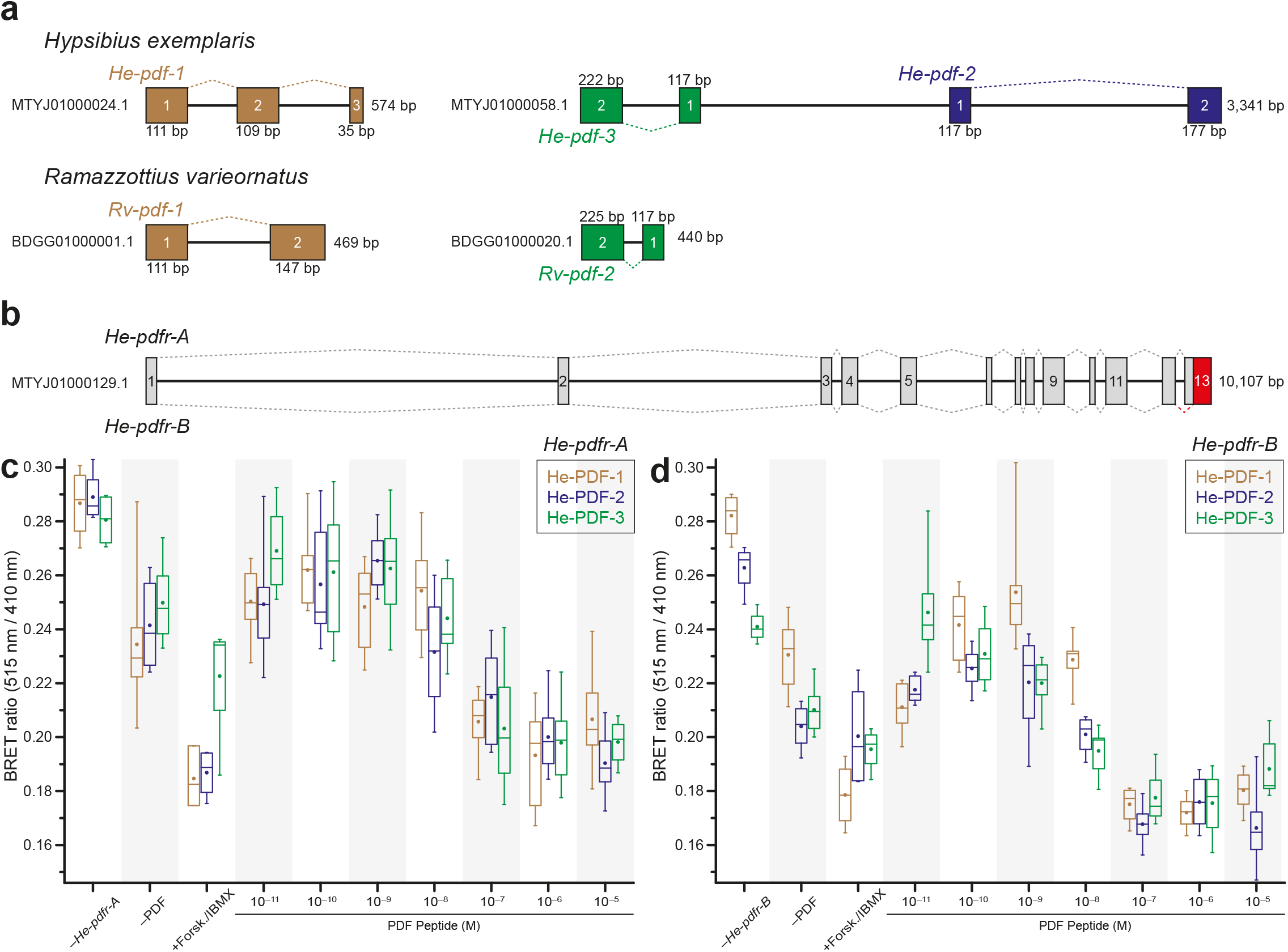
Genomic structure of *pdf* and *pdfr* genes of eutardigrades, and results of *in vitro* functional analyses of PDFs and PDFR isoforms in *H. exemplaris*. (**a, b**) Diagrams of genomic scaffolds illustrating location and orientation of *pdf* and *pdfr* genes in *H. exemplaris* and *Ramazzottius varieornatus*. Exons are depicted as colored rectangles. Dashed lines indicate position of introns. Note proximity of *He-pdf-2* and *He-pdf-3* on single scaffold (MTYJ01000058.1). Red rectangle (exon 13 of *He*-*pdfr*-*B* in **b**) indicates alternative 3’ splice junction (acceptor site) within last exon of *He*-*pdfr* in *H. exemplaris*. (**c**, **d**) BRET ratio changes of HEK293T cells transfected with long (*He-pdfr-A*) and short variants (*He-pdfr-B*) of PDF receptor from *H. exemplaris* after activation with synthetic PDFs (He-PDF-1, He- PDF-2 and He-PDF-3) in dose-dependent manner. Note decreasing BRET ratios (i.e., increasing intracellular cAMP levels) at higher concentrations of peptide indicating stronger response to stimuli. Box plots of eight biological replicates (n=8). Boxes correspond to interquartile ranges. Circle inside each box represents arithmetic mean. Negative controls were performed without *pdfr* transfection (*−He-pdfr-A*; *−He-pdfr-B*) and PDF stimuli (−PDF). Positive control was conducted with Forskolin/IBMX (+Forsk./IBMX), which causes high intracellular concentrations of cAMP.

### One gene but two isoforms of PDF receptor in *H. exemplaris*

After the initial BLAST screening, complete coding sequence of the putative homolog of *pdf receptor* (*pdfr*) gene of *H. exemplaris*, *He-pdfr*, was obtained from each of the three publicly available transcriptome assemblies (GenBank accession numbers: GBZR01000000, GFGW01000000, and GJGU01000000). The existence of a single *pdfr* gene in *H. exemplaris* was confirmed by genomic searches (MTYJ00000000.1). The respective gene is 10,107 bp long and located on the reverse strand of scaffold0129 (MTYJ01000129.1). It consists of thirteen exons and twelve introns (Fig. 2b; Supplementary Data 1). Cloning and sequencing, however, revealed two transcribed isoforms, suggesting that at least two splice variants of PDFR are present in *H. exemplaris*, *He*-*pdfr*-*A* (1,581 bp) and *He*-*pdfr*-*B* (1,557 bp), the latter being eight amino acids (24 bp) shorter in the c-terminal region due to an alternative 3’ splice junction (acceptor site) in the last exon of *He-pdfr* (Fig. 2b; Supplementary Data 1). The orthologous gene *Rv-pdfr* of *Ramazzottius varieornatus* is considerably shorter (5,341 bp) than *He-pdfr* of *H. exemplaris* and consists of twelve rather than thirteen exons. Like *He- pdfr*, *Rv-pdfr* is located on the reverse strand of the respective scaffold (scaffold0005: BDGG01000005.1) and the *Rv-pdfr* transcript is of comparable length (1,554 bp) to those of *He-pdfr-A* and *He-pdfr-B* (Supplementary Data 1).

Using a similar strategy, we further identified two putative *pdfr* homologs in the transcriptomic and/or genomic databases from the eutardigrade *Paramacrobiotus metropolitanus* and one putative *pdfr* homolog in the heterotardigrades *Echiniscus testudo*, *Echiniscoides sigismundi*, and *Batillipes* sp. To clarify the orthology of detected homologs, we performed a phylogenetic analysis of members of class B G protein-coupled receptors across bilaterians. In the resulting maximum likelihood tree, PDFRs occur as the sister clade to the corticotropin-releasing hormone receptors (Fig. 1c; Supplementary Fig. 2), thus confirming the identity of putative homologs from tardigrades, including both splice variants from *H. exemplaris*, as members of the *pdfr* clade.

### Functionality of PDFs and PDFR isoforms in *H. exemplaris*

To test the functionality of the two splice variants of PDFR and the three PDFs of *H. exemplaris*, we established Bioluminescence Resonance Energy Transfer (BRET) assays monitoring changes in the intracellular cAMP concentration upon PDF stimulus. Different dosages of synthetic He-PDF-1, He-PDF-2 and He-PDF-3 at final concentrations ranging from 10^−1^^1^ M to 10^−5^ M (n=8 for each dosage) induced PDFR/cAMP responses in the respective cell cultures (Fig. 2c, d; Supplementary Data 1). Interestingly, the resulting box plots for both receptors are similar across the three peptides. This suggests that both receptors, He-PDFR-A and He-PDFR-B, recognize all three peptides: He-PDF-1, He-PDF-2, and He-PDF-3.

### Localization of PDFs and *pdf* mRNA in *H. exemplaris*

We used newly generated antibodies to localize the tardigrade PDFs. To test the specificity and potential cross reactivity of these antibodies as well as the previously used β-PDH serum^12^, we performed western blots with synthetic He-PDF-1, He-PDF-2, and He-PDF-3 (Supplementary Fig. 3a–d). For control experiments, the three specific PDF antibodies were applied reciprocally to the individual peptides in each blot. The results show that each of the three antibodies recognizes the corresponding peptide, indicated by a distinct band slightly below 4.6 kDa, while not binding to the non-corresponding PDFs (asterisks in Supplementary Fig. 3a–c). Like in the controls, the β-PDH serum does not show any staining (Supplementary Fig. 3d), suggesting that it does not cross-react with any of the three PDFs, which is consistent with the previous negative result^12^.

Immunolocalization using specific antibodies revealed signal in several regions of the central nervous system and two extracerebral cells (Figs 3–5). The most prominent signal occurs in somata of two bilateral pairs of dorsal unipolar neurons (inner lobe cells) located in the posterior region of the inner lobes of the brain (Fig. 3a, e, f; Supplementary Movie 1). Each pair of these neurons sends out anteriorly their axons towards the contralateral region of the brain (Fig. 3a, f). The axons of each pair then fasciculate with those of their contralateral counterpart while passing through the anterior region of the central brain neuropil to the other brain hemisphere (Fig. 3f; Supplementary Movie 1). The two axonal pairs separate from each other and loop posteriorly descending towards the first trunk ganglion, to which they pass via the inner connectives (Fig. 3f). Each pair then passes through the contralateral side of each of the first three trunk ganglia, i.e., the two axons from the right brain hemisphere pass through the left side of each trunk ganglion and vice versa (Fig. 4a–p). The axons of the inner lobe cells terminate in two pairs of button-like structures in the antero-ventral region of the fourth trunk ganglion (arrows in Fig. 4e, j, o).

**Fig. 3.**
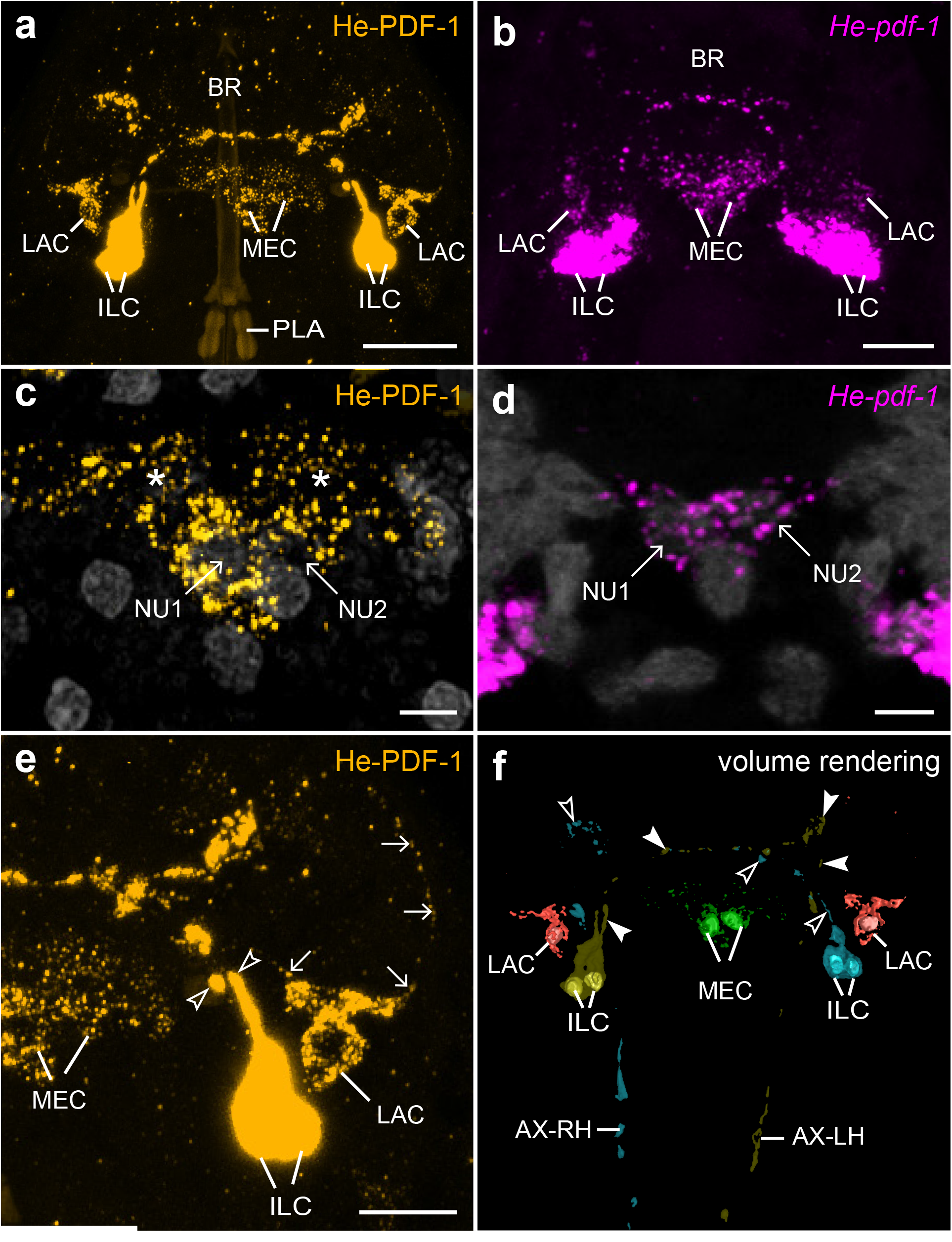
Localization of He-PDF-1 and *He-pdf-1* transcripts in the brain of the tardigrade *H. exemplaris*. Antibody (a, c, e) and mRNA labeling (b, d). 3D reconstruction (f) and projections of confocal substacks (a–e) in dorsal view. Anterior is up in all images. Pharyngeal placoids are autofluorescent in a. DNA staining is illustrated in gray in c and d. (a) Localization of He-PDF-1 peptide in inner lobe cells, lateral cells, and median cells. (b) Localization of *He-pdf-1* mRNA. Note correspondence to immunolabeling in a. (c) Detail of immunolabeled median cells with their nuclei (NU1 and NU2). Asterisks indicate varicosities of neurites labeled with He-PDF-1 antibody. (d) Detail of *He-pdf-1* mRNA expression in median cells. Note correspondence to immunolabeling in c, except that varicosities have not been labeled. (e) Detail of right brain hemisphere from specimen in a showing individual axons of inner lobe cells (open arrowheads) and neurites of lateral cell (arrows). (f) Volume rendering of He-PDF-1 immunoreactivity illustrating number and position of somata of individual neurons with nuclei (colored). Note immunoreactivity in two pairs of inner lobe cells (yellow and cyan), two median cells (green), and two lateral cells (red) withinb the brain. Arrowheads indicate varicosities of axonal projections of left (filled arrowheads) and right pairs of inner lobe cells (open arrowheads) that cross over to other brain hemisphere. Abbreviations: AX-LH axonal projections from left brain hemisphere, AX-RH axonal projections from right brain hemisphere, BR brain, ILC inner lobe PDF-immunoreactive cells, LAC lateral PDF-immunoreactive cells, MEC median PDF-immunoreactive cells, NU1 nucleus of median cell 1, NU2 nucleus of median cell 2, PLA pharyngeal placoids. Scale bars: 10 µm (a), 5 µm (b, e), and 2 µm (c, d).

**Fig. 4.**
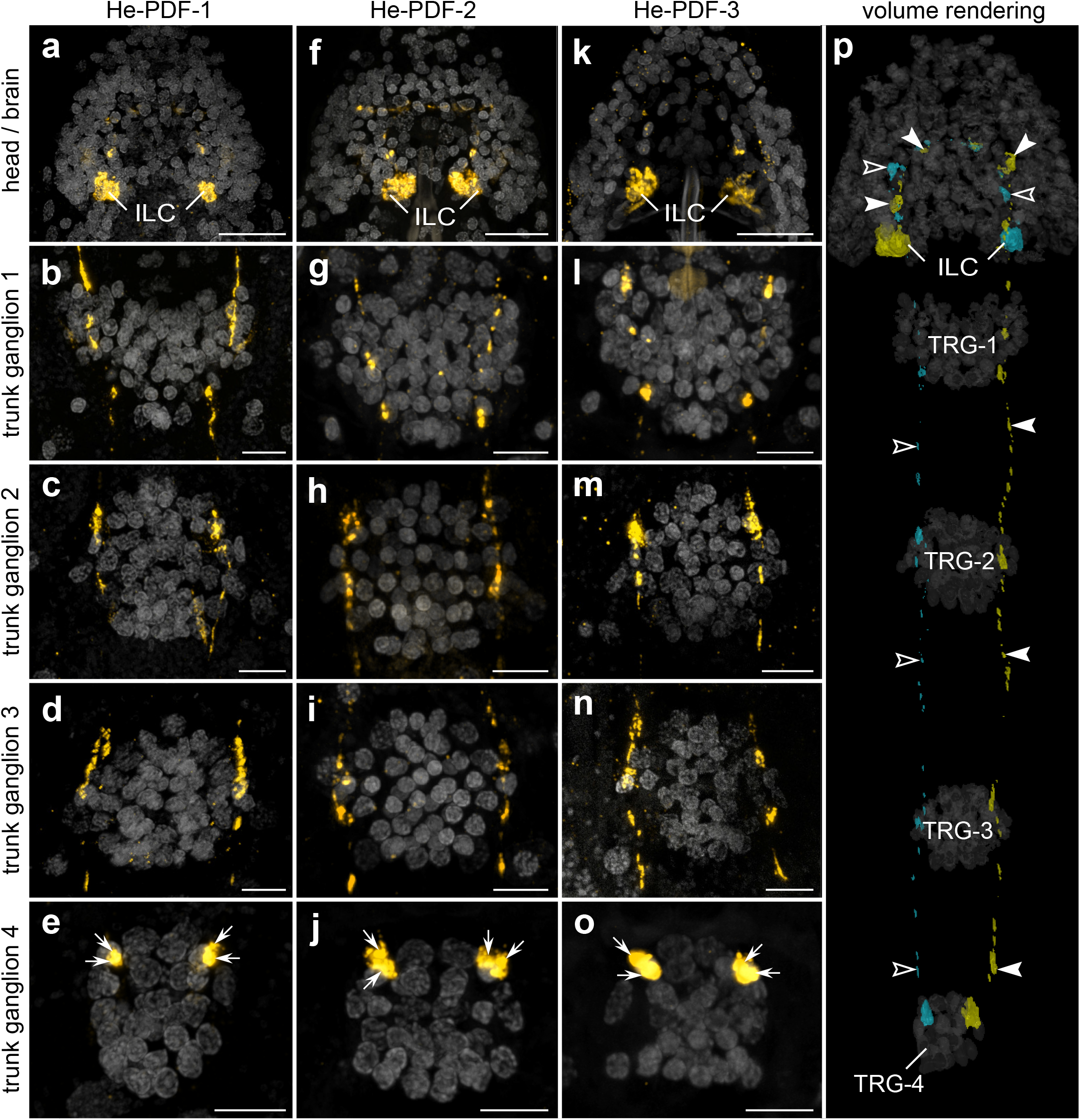
Localization of each of the three PDFs in the nervous system of *H. exemplaris*. Antibody labeling (orange in **a–o**), and 3D reconstruction (yellow and cyan in **p**). DNA staining is illustrated in gray. Projections of confocal substacks of head region in dorsal view (**a, f, k**) and trunk ganglia in ventral view (**b–e, g–j, l–o).** Anterior is up in all images. (**a–o**) Note prominent expression of all three peptides in two pairs of somata in inner lobes of brain. Varicose axonal projections pass through first three trunk ganglia and terminate in anterior region of fourth trunk ganglion (arrows in **e, j, o**). (**p**) Volume rendering of left (yellow) and right pairs (cyan) of inner lobe cells and trajectories of their contralateral fibers (filled and open arrowheads) based on He-PDF-1 immunolabeling. Dorsal perspective. Note position of potential terminal buttons in anterior region of fourth trunk ganglion. Abbreviations: ILC inner lobe PDF-immunoreactive cells, TRG-1**–**TRG-4 first to fourth trunk ganglia. Scale bars: 10 µm (**a, f, k**), and 5 µm (**b–e, g–j, l–o**).

**Fig. 5.**
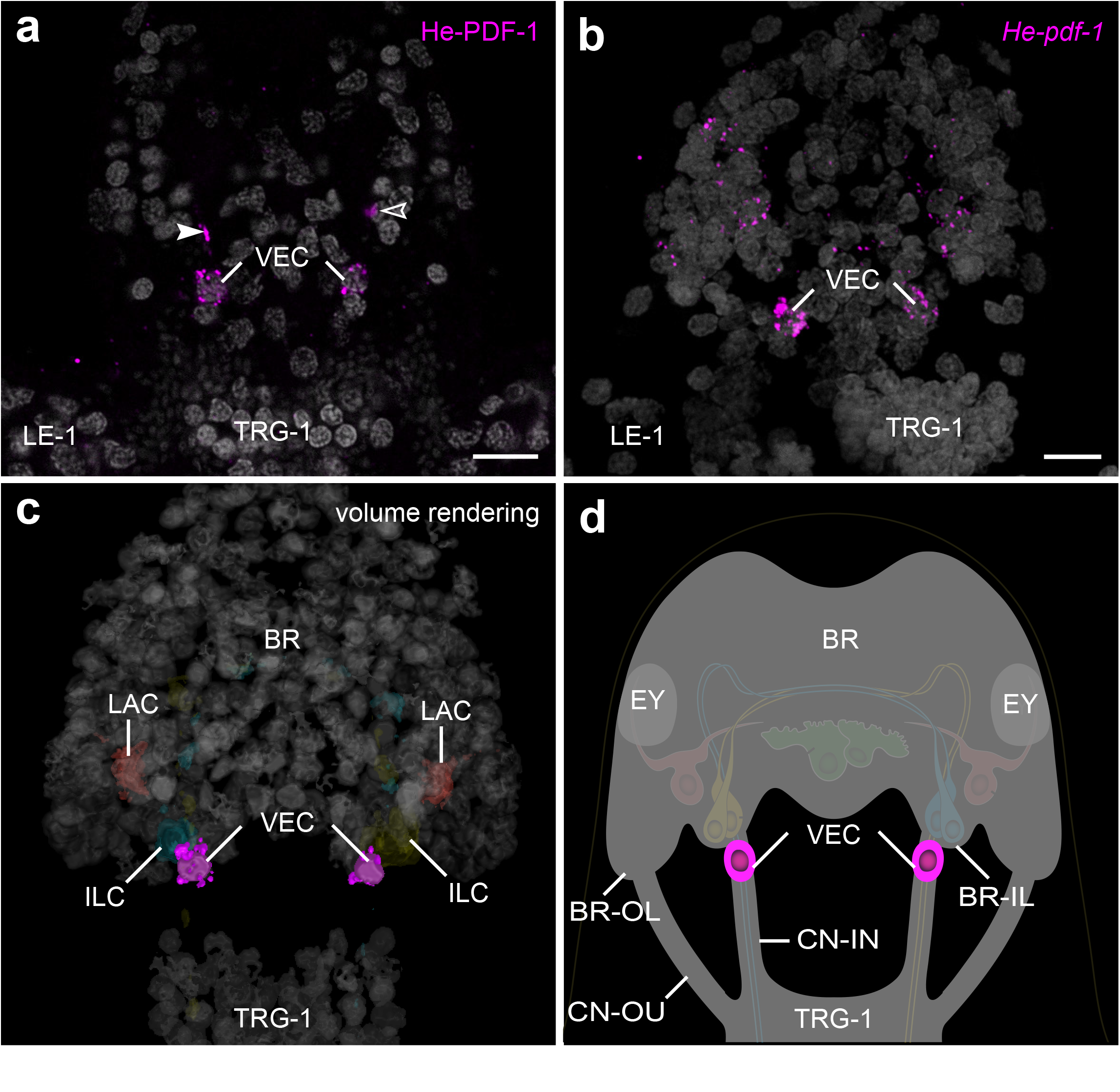
Localization of He-PDF-1 and *He-pdf-1* transcripts in the ventral extracerebral PDF-*ir* cells of *H. exemplaris*. Specimens are presented in ventral perspective. Anterior is up. DNA staining is illustrated in gray in **a–c**. (**a, b**) Projections of confocal substacks. (**a**) Antibody labeling of He-PDF-1 peptide. Arrowheads indicate immunoreactive fibers of inner lobe cells in inner connectives. (**b**) Localization of *He-pdf-1* mRNA transcripts, which corresponds to distribution of He-PDF-1 peptide in **a**. (**c**) Volume rendering of immunolabeled anterior body region. Note position of somata of both ventral extracerebral PDF-*ir* cells outside the brain, anterior to first trunk ganglion. (**d**) Diagram of anterior body region illustrating position of ventral extracerebral PDF-*ir* cells that are widely separated from other PDF-*ir* cells across the z axis. Abbreviations: BR brain, BR-IL inner lobe of brain, BR-OL outer lobe of brain, CN-IN inner connectives, CN-OU outer connectives, EY eyes, ILC inner lobe PDF-immunoreactive cells, LAC lateral PDF-immunoreactive cells, LE-1 first leg, TRG-1 first trunk ganglion, VEC ventral extracerebral PDF-immunoreactive cells. Scale bars: 5 µm (**b, c**).

Besides one pair of inner lobe cells, each brain hemisphere of *H. exemplaris* exhibits one lateral and one median cell, totaling eight He-PDF-1-*ir* neurons in the entire brain (Fig. 3a, c, e, f). The He-PDF-1*-ir* signal in the two median cells indicates anaxonic morphology, with numerous neurites in the anterior region of each cell but no apparent axon (Fig. 3a, c, e, f; Supplementary Movie 1). In contrast to the median cells, each lateral cell shows typical pseudounipolar structure, with a neurite arising from the soma that bifurcates into a median and a lateral process (arrows in Fig. 3a, e). We could not trace the entire path of the median neurite of the lateral cell, but it seems to enter the central neuropil, where most neurites of the median cells are located. The lateral neurite of the lateral cell shows an arcuate shape (arrows on the right of Fig. 3e). It terminates at the dorsal surface of the rhabdomeric cell of the eye, which is situated in the outer lobe of the brain (Supplementary Fig. 4). While the four inner lobe cells and their axons are immunoreactive to all three peptides (Fig. 4a–o; Supplementary Fig. 5), only He-PDF-1-*ir* signal (detectable at higher laser intensities) occurs in the two median and the two lateral cells, irrespective of the Zeitgeber time (Supplementary Fig. 6a– c).

In addition to the eight He-PDF-1-*ir* somata within the brain, there are He-PDF-1-*ir* somata outside the brain that belong to two individual cells. These ventral extracerebral cells are situated between the brain and the first trunk ganglion (Fig. 5a–d; Supplementary Movie 2). They are ventrally adjacent to the inner connectives of the central nervous system but do not show any PDF-*ir* fibers themselves. The PDF immunoreactivity in the inner connectives (arrowheads in Fig. 5a) solely derives from axons and varicosities of the inner lobe cells. Like the median and lateral cells within the brain, the ventral extracerebral cells show relatively weak He-PDF-1*-ir* signal and no anti-He-PDF-2 or anti-He-PDF-3 immunoreactivity.

To further substantiate the results of immunolabeling, we performed various sets of *in situ* hybridization experiments. The mRNA labeling essentially revealed the same results, except that neuronal projections are labeled weaker using this technique as compared to the immunolabeling. All three complementary probes to the *He-pdf-1*, *He-pdf-2* and *He-pdf-3* transcripts are co-localized in the inner lobe cells, whereas the median cells, the lateral cells and the ventral extracerebral cells exhibit only *He-pdf-1* signal (Figs 3b, d, 5b; Supplementary Movies 3, 4). This finding is consistent in single and double *in situ* hybridization experiments (Supplementary Figs 7, 8) as well as assays using a combination of the *He-pdf-1* probe with the He-PDF-1 antibody (Supplementary Fig. 9). Like the immunostaining experiments, mRNA labeling in specimens at different Zeitgeber times revealed the most prominent *He-pdf-1* signal in the somata of the inner lobe cells, whereas the median and lateral cells show relatively weak staining (Supplementary Fig. 6d–f). *He-pdf- 1* signal is consistently stronger in somata of the inner lobe cells (n=15), whereas *He-pdf-2* (n=5), and especially *He-pdf-3* (n=5) show relatively weaker signals, even at higher concentrations of the respective probes (Supplementary Table 6) and laser intensities used (Fig. 6a–c). The same holds true for the results of double labeling with each two of the three probes (Supplementary Fig. 8). This noticeable difference in relative signal intensities corresponds to the numbers of short sequencer reads we quantified from the available transcriptomes using a read mapping approach, which revealed nearly a tenfold lower expression of *He-pdf-2* and a hundredfold lower expression of *He-pdf-3* compared to *He*-*pdf*- *1* (Fig. 6d, e; Supplementary Data 2).

**Fig. 6.**
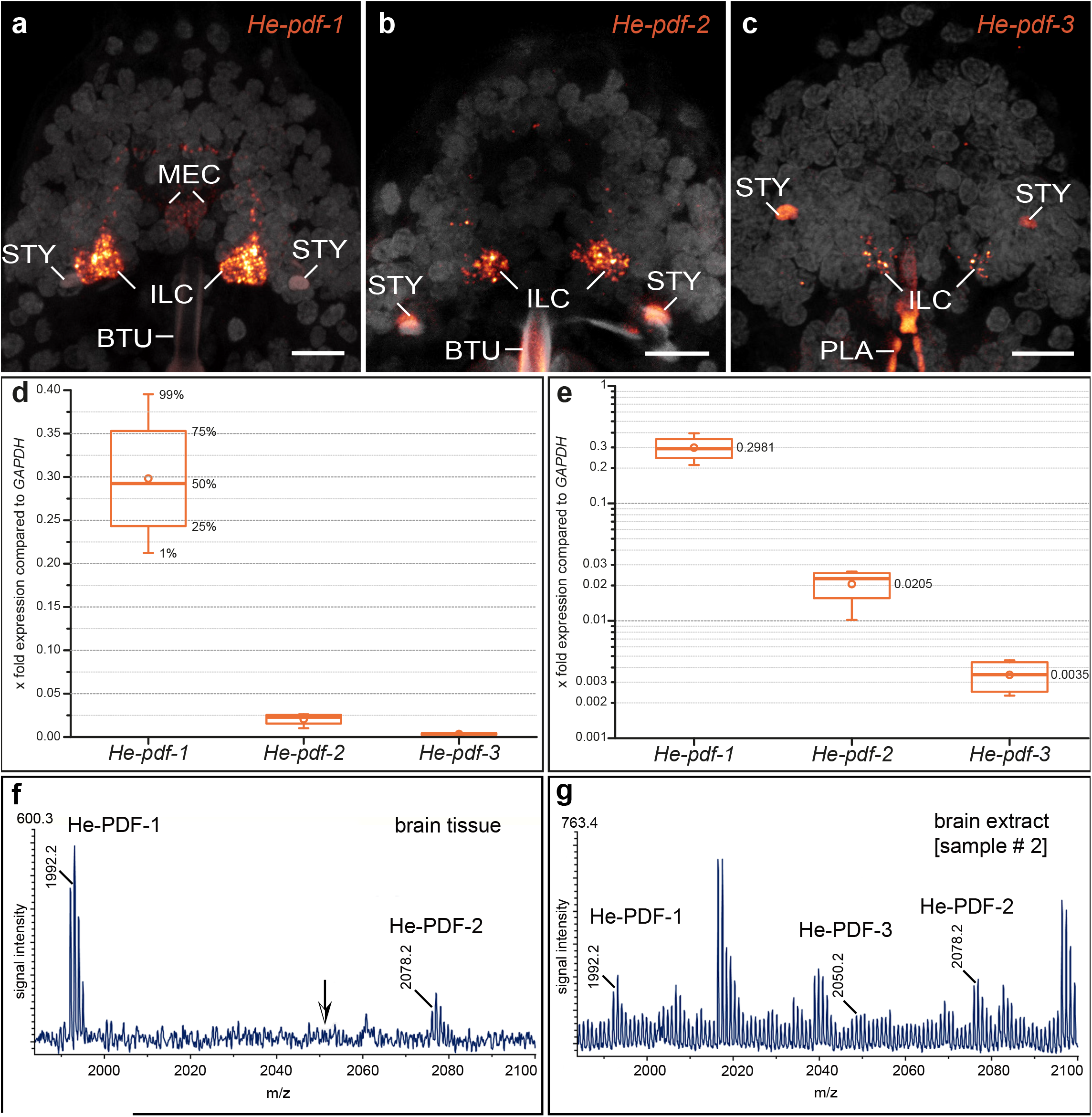
Differences in expression levels of the three *pdf* genes and MALDI-TOF mass spectra of the three PDFs in *H. exemplaris*. (**a–c**) Detection of *He-pdf-1*, *He-pdf-2* and *He- pdf-3* mRNA using same settings of confocal microscope. Brains in dorsal view. Anterior is up. DNA staining is shown in gray. Buccal tube, stylets, and pharyngeal placoids are autofluorescent. Note that *He-pdf-1* shows highest and *He-pdf-3* lowest signal in inner lobe cells. (**d, e**) Relative expression levels of *He-pdf-1*, *He-pdf-2* and *He-pdf-3* based on comparative analysis of raw sequence reads from four independent RNA-seq experiments using short read mapping approach, with housekeeping gene *GAPDH* as reference. Boxes correspond to interquartile ranges. Circle inside each box represents arithmetic mean. (**d**) Linear chart. (**e**) Logarithmic chart using same dataset as in **d**. (**f, g**) Representative MALDI- TOF mass spectra obtained from single dissected brain by direct tissue profiling (**f**) and extract of 100 brains (**g**) from sample #2 (see Methods; Supplementary Fig. 10). Ion signals are marked and represent single charged peptides [M+H]^+^. Note that using direct tissue profiling only He-PDF-1 and He-PDF-2 could be detected, whereas ion masses corresponding to all three predicted He-PDFs were obtained from brain extracts. Abbreviations: BTU buccal tube, ILC inner lobe PDF-immunoreactive cells, MEC median PDF-immunoreactive cells, PLA pharyngeal placoids, STY stylet. Scale bars: 5 µm (**a–c**).

### Identification of PDFs in brain samples from *H. exemplaris* by mass spectrometry

To confirm the presence of putative PDFs in the brain, we first analyzed samples of entire brains using direct tissue profiling by matrix-assisted laser desorption/ionization (MALDI) time of flight (TOF) mass spectrometry (MS). This revealed a typical spectrum in a mass range at m/z 1,800–2,100 Da (Fig. 6f, g). The most abundant ion signal originated from He- PDF-1, followed by He-PDF-2, whereas the predicted ion mass of He-PDF-3 could not be detected (arrow in Fig. 6f). Further fragmentation experiments using MALDI-TOF/TOF MS to determine the amino acid sequences were unsuccessful due to the low concentration of peptide in the sample spot. We then prepared brain extracts to increase the concentration of peptides in the sample sets. The resulting MALDI-TOF mass spectrum revealed all three predicted PDFs, where He-PDF-1 shows the most abundant ion signal, followed by He-PDF- 2 and He-PDF-3 (Fig. 6g). This corresponds well to the results of mRNA detection (Figs 3e, 6a–c) as well as quantitative analyses of expression levels (Fig. 6d, e). For chemical identification of transcriptome-predicted PDFs, we analyzed brain extracts by ESI-Q Exactive Orbitrap MS, followed by an evaluation of peptide fragmentation using PEAKS 10.5 software package. This resulted in a confirmation of amino acid sequences of all three PDFs (Supplementary Fig. 10).

### Quantification and localization of PDF receptor in *H. exemplaris*

Quantitative analysis of four transcriptomes of *H. exemplaris* revealed similar expression levels of both splice variants of the PDFR, *He-pdfr-A* and *He-pdfr-B*, each of which is tenfold lower than that of *He-pdf-1* (Supplementary Fig. 11; Supplementary Data 2). To localize cells that express *pdfr*, we conducted Hybridization Chain Reaction – Fluorescence *In Situ* Hybridization (HCR-FISH) using DNA split initiator *He-pdfr* probes. Due to the small difference of 24 nucleotides, we could not distinguish between the two isoforms of the receptor. Our data show a widespread expression of *pdfr* in all major organs and tissues of *H. exemplaris*, including the eye, the outer, inner, and median lobes of the brain, the four trunk ganglia, the somatic muscles, the digestive tract (pharynx, esophagus, and midgut), the trophocytes of the ovary, the Malpighian tubules, the stylet glands, and the claw glands (Fig. 7a–k; Supplementary Fig. 12). We further localized *He-pdfr* mRNA in storage cells that are abundant in the body cavity (Fig. 7l). Double HCR-FISH revealed co-expression of *He-pdfr* with visual *r-opsin* (*He-r-opsin-v*) in the rhabdomeric cell of the eye (Fig. 7a; Supplementary Fig. 13). The same technique exhibited *He-pdfr* mRNA in somata of all PDF-*ir* neurons within the brain, including the inner lobe cells, the lateral cells, and the median cells (Fig. 7c; Supplementary Fig. 14). The results of positive control experiments using two sets of redundant probes are consistent and confirm the specificity of our *He*-*pdfr* probes (Supplementary Fig. 15).

**Fig. 7.**
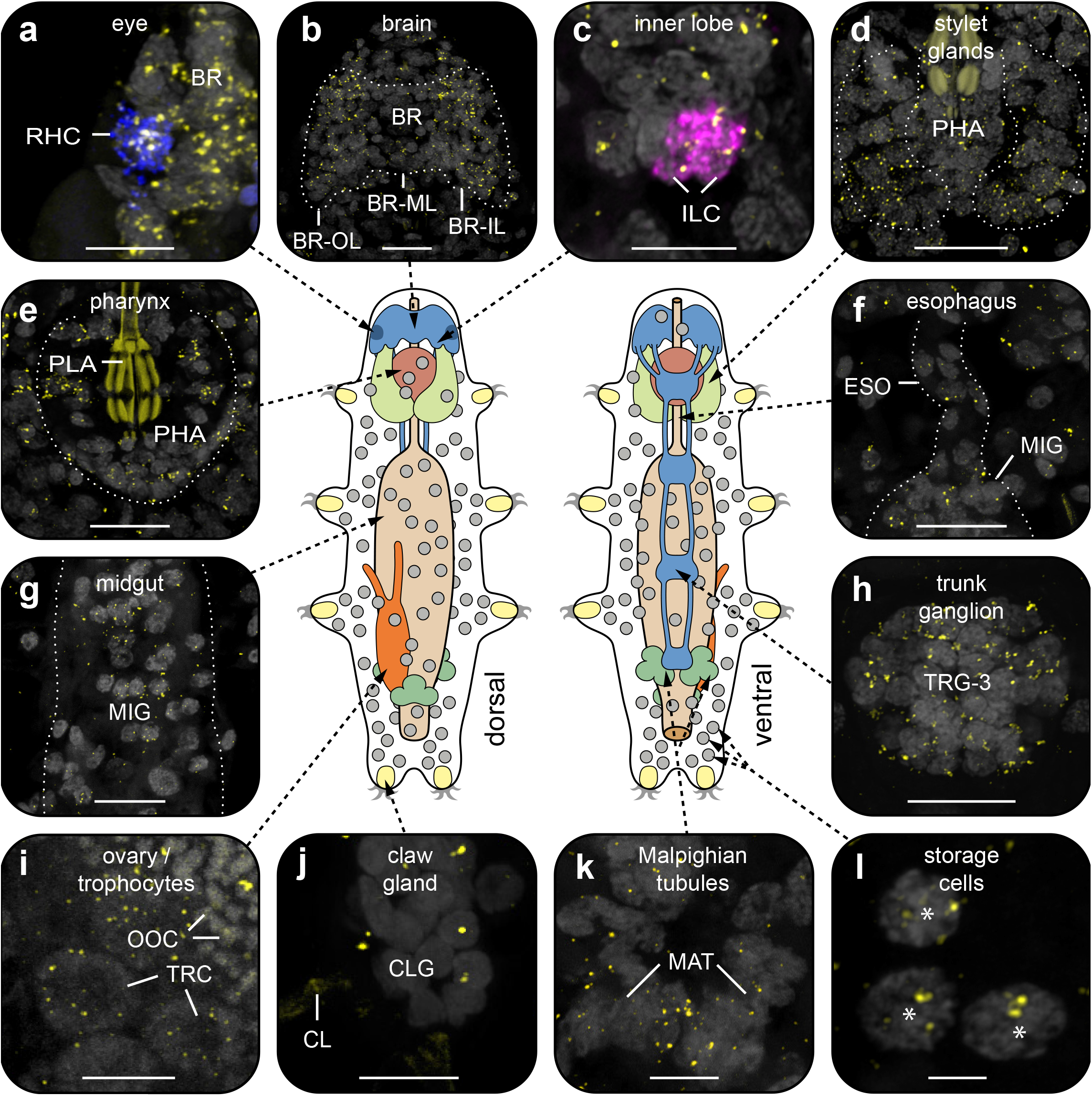
Localization of PDF receptor mRNA in various tissues and cells of *H. exemplaris*. Labeling using *He-pdfr* probes is illustrated in yellow in **a–l** (see Supplementary Fig. 15 for positive controls). Projections of confocal substacks (**a–l**) and diagrams of tardigrade anatomy in dorsal and ventral views (center). Dotted white lines indicate contours of organs and tissues. Anterior is up in all images. Gray color shows DNA staining in **a–h, j,** and **k**, and autofluorescence in **i** and **l**. Pharyngeal placoids are autofluorescent in **d** and **e**, and claws in **j**. (**a**) Expression of *He*-*pdfr* in rhabdomeric cell of eye labeled with *r-opsin-v* mRNA probe (blue). (**b**) Expression of *He-pdfr* in the brain. (**c**) Expression of *He*-*pdfr* in inner lobe cells double labeled with *He-pdf-1* probe (magenta). (**d–l**) Expression of *He-pdfr* in various other tissues and cells. Asterisks indicate storage cells that occur within body cavity. Abbreviations: BR brain, BR-IL inner lobe of brain, BR-ML median lobe of brain, BR-OL outer lobe of brain, CLG claw gland, CL claw (visualized by cuticular autofluorescence), ESO esophagus, ILC inner lobe PDF-immunoreactive cells, MAT Malpighian tubules, MIG midgut, OOC autofluorescent yolk granules of an oocyte, PHA pharynx, PLA pharyngeal placoids, RHC rhabdomeric cell, TRG-3 third trunk ganglion, TRC trophocytes. Scale bars: 10 µm (**b**, **d**, **e–h**), 5 µm (**a**, **c**, **i–k**), and 2 µm (**l**).

## Discussion

### Scenario on the evolution and duplication of *pdf* genes in tardigrades

Comparison with previous findings^12^ allows us to expand the scenario on the evolution of *pdf* genes (Fig. 1a). While only one *pdf* gene was most likely present in the last common ancestor of protostomes, a duplication might have led to two genes in the ecdysozoan lineage, *pdf-I* and *pdf-II*, whose homologs are still present in extant priapulids, nematodes, and onychophorans^11,12^. Subsequently, *pdf-II* was lost whereas *pdf-I* has been retained in tardigrades and arthropods, followed by independent duplications in each of these taxa^12^. While *pdf* genes have not been characterized in heterotardigrades, the last common ancestor of eutardigrades likely possessed two *pdf-I* homologs, one of which was duplicated either once again in the parachelan lineage, followed by subsequent losses in some species (*Ramazzottius varieornatus* and *Mesobiotus philippinicus*), or multiple times independently in separate lineages, including those containing *Richtersius coronifer*, *H. exemplaris*, and *Paramacrobiotus* species.

Although a conclusive scenario on the evolution of *pdf* genes is hard to depict due to the still unresolved internal phylogeny of Parachela^39^, a single duplication of the parachelan *pdf-2/3* precursor, followed by subsequent species- or lineage-specific losses of either *pdf-2* or *pdf-3*, is in line with the number of identified *pdf* genes in different eutardigrade species. This scenario received additional support from the topology of our maximum likelihood tree, in which parachelan PDFs occur in two distinct clades (PDF-1 and PDF-2/3). Despite the clear indication of two *pdf-I* paralogs (*pdf-1* and *pdf-2/3* precursor) in the last common ancestor of eutardigrades, we caution that this scenario is still based on a limited number of sequenced tardigrade genomes and relatively short peptide sequences, consisting only of 18 amino acids. Hence, our hypothesis on the evolution and duplication of *pdf* genes in tardigrades will have to be further tested once additional genomes, especially from heterotardigrades and apochelan eutardigrades, have become available.

### Homology of protocerebral PDF-*ir* neurons and the origin of ventral extracerebral PDF-*ir* cells in *H. exemplaris*

We have demonstrated that the three *pdf* homologs of *H. exemplaris*, which are derivatives of *pdf-I* of the last common ancestor of Ecdysozoa^12^ (Fig. 1a), exhibit different expression patterns. While *He*-*pdf-1*, *He*-*pdf-2* and *He*-*pdf-3* are co-expressed in the two pairs of inner lobe cells, only *He*-*pdf-1* mRNA is found in the median and lateral cells as well as the ventral extracerebral PDF*-ir* cells (Fig. 8a–c). The results of mRNA labeling are entirely consistent with those based on the immunolocalization of PDFs, demonstrating that eight out of ten PDF*-ir* somata of *H. exemplaris* are located in the brain. Given that the tardigrade brain most likely consists of a single segmental region homologous with the protocerebrum of other panarthropods^40^, the existence of protocerebral He-PDF-*ir* neurons corresponds to that in onychophorans^11,12^ and most arthropods^17,24,41^. This suggests that several bilateral PDF-*ir* somata were present in the protocerebrum of the last common ancestor of Panarthropoda. Provided that the median protocerebral PDF-*ir* cells are homologous across panarthropods, their absence in *Drosophila melanogaster*, *Rhyparobia madeira*, *Apis mellifera*, *Acyrthosiphon pisum*^5,9,21,42^ and various other insects^4,13^ most likely resulted from convergent losses in the corresponding taxa.

**Fig. 8.**
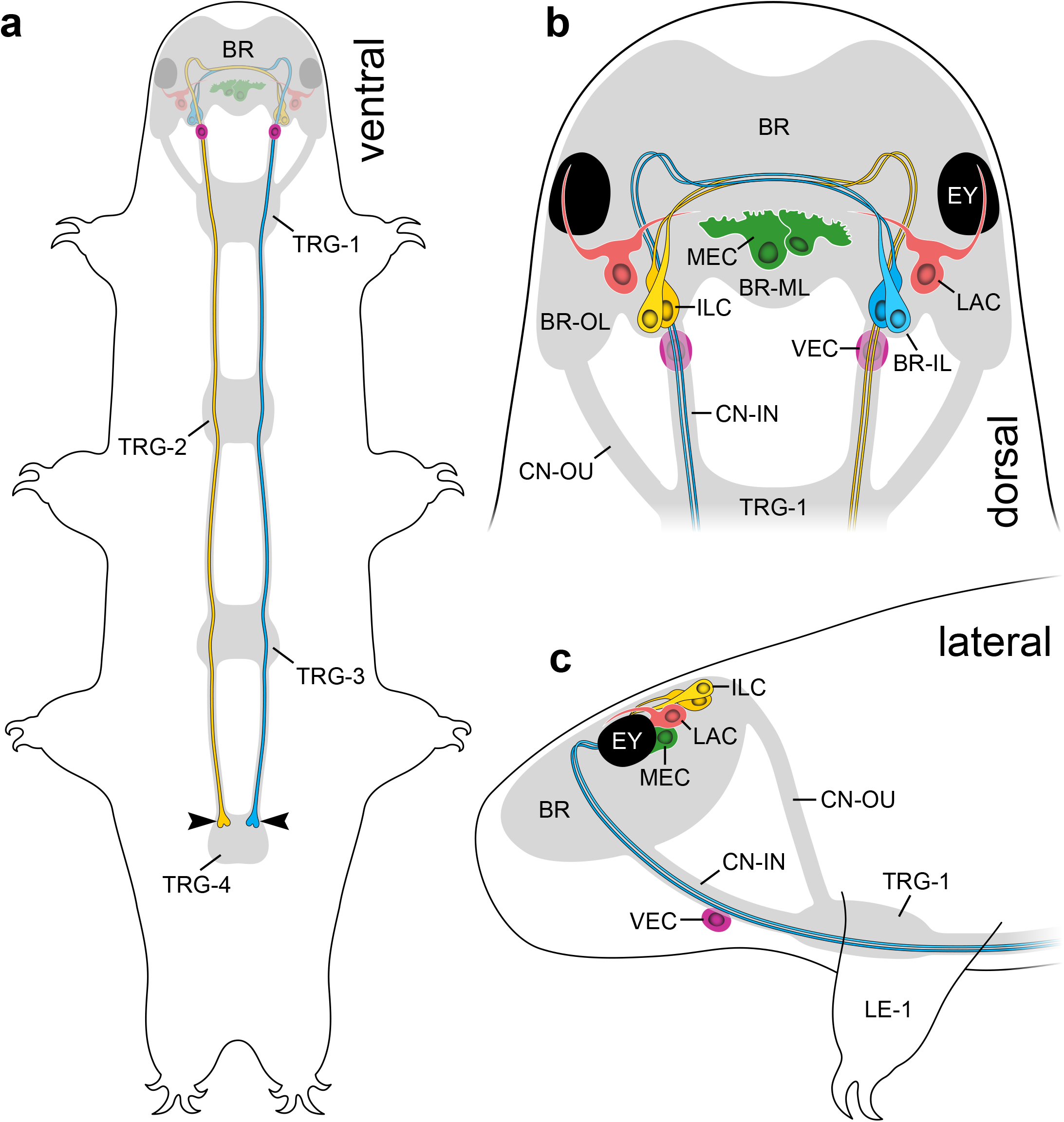
**Summary diagrams of localization of PDFs in the head of *H. exemplaris***. Brain, trunk ganglia and connectives are indicated in gray. (**a**) Overview illustrating position of PDF-*ir* neurons (colored), ventral extracerebral PDF-*ir* cells, and trajectories of inner lobe cells (yellow and blue). Arrowheads point to button-like terminals on antero-ventral surface of fourth trunk ganglion. (**b**) Diagram illustrating number, position, morphology, and connectivity of PDF-*ir* cells in the head in dorsal view. Note eight cerebral neurons exhibiting different morphologies and two ventral extracerebral cells associated with inner connectives. Note also that lateral PDF-*ir* neurons send their projections to eyes and central part of brain. (**c**) Same as in **b**, but in lateral view. Abbreviations: BR brain, BR-IL inner lobe of brain, BR-ML median lobe of brain, BR-OL outer lobe of brain, CN-IN inner connectives, CN-OU outer connectives, EY eye, ILC inner lobe PDF-immunoreactive cells, LAC lateral PDF-immunoreactive cells, LE-1 first leg, MEC median PDF-immunoreactive cells, TRG-1**–**TRG-4 first to fourth trunk ganglia, VEC ventral extracerebral PDF- immunoreactive cells.

The two ventral extracerebral PDF-*ir* cells associated with the inner connectives of *H. exemplaris* (Fig. 8b, c) are most likely a derived feature of tardigrades or a tardigrade subclade, as such extraganglionic cells situated between the brain and the ventral nervous system have not been reported from any other animal group. These cells might correspond to a pair of “anteroventral cells” that arise early in the embryo of *H. exemplaris* (formerly referred to as *H. dujardini*^33^) and become connected to the first trunk ganglion as well as the circumbuccal nerve ring^43^. These embryonic cells give rise to neurites extending towards the body surface later in development, suggesting they are sensory neurons. Our labeling did not reveal any PDF-*ir* fibers associated with the ventral extracerebral cells, which might be due to the relatively weak staining of these cells. Thus, the identity, detailed morphology, and potential function of the ventral extracerebral PDF-*ir* cells remain to be clarified.

### PDFs of tardigrades are neurohormones and neuromodulators that control various body functions

The results of our *in vitro* assays using BRET in transfected human cells indicate that all three PDFs of *H. exemplaris* and both splice variants of their receptor are functional and that G protein-coupled receptor signaling (in this case PDF/PDFR signaling) based on cAMP regulation is present in tardigrades. We further found no conspicuous differences in the cAMP response patterns, neither between the three peptides nor the two splice variants of their receptor, which contrasts with the apparent dose-dependent differences between the individual PDFs and their isoforms in the nematode *Caenorhabditis elegans*^29^, the onychophoran *Euperipatoides rowelli*^11^, and the crustacean *Carcinus maenas*^18^. However, our expression level analyses revealed substantial differences in the abundance of the three peptides in *H. exemplaris*, with *He-pdf-2* showing tenfold lower and *He-pdf-3* hundredfold lower values than *He-pdf-1*. The functional role of these differences between the three peptides as well as their different localization in the head (Fig. 8a–c) is unclear, given that we did not observe noticeable differences in their interaction with either splice variant of PDFR in transfected cells. However, they might still be subject to distinct regulatory mechanisms *in vivo*, i.e., in the tardigrade body, which we did not investigate.

Our data further revealed that PDFR of *H. exemplaris* is expressed in all major organ systems and tissues of the body, including storage cells. Co-localization of PDFR with all three PDFs in somata of the inner lobe cells and with He-PDF-1 in the median and lateral cells suggests an autoregulatory signaling loop in tardigrades. This might resemble the PDF/PDFR autoreception in the insect clock^35,44,45^, although PDF and PDFR are co-localized only in a subset of PDF-*ir* clock neurons in insects^36,37^. Analyzing the expression of core and associated clock genes in *H. exemplaris* may help to clarify whether PDF-*ir* neurons are involved in the circadian system of tardigrades.

The spatial separation of PDF-producing neurons and *pdfr-*expressing cells suggests that the PDFs of tardigrades act as neurohormones that might be released into the body cavity for targeting other cells. We have indeed detected two pairs of prominent button-like structures on the antero-ventral surface of the fourth trunk ganglion (arrowheads in Fig. 8a) that represent the axonal terminals of the inner lobe cells, the somata of which are located in the brain. These paired, unipolar neurons project axons to the contralateral brain hemisphere, which then follow each inner connective and pass through the first three trunk ganglia to finally terminate in the fourth trunk ganglion (Fig. 8a–c). We speculate that these button-like structures are potential release sites of all three peptides, as the inner lobe cells are the only neurons that produce all three of them. Thus, the inner lobe cells might play a neuroendocrine role in *H. exemplaris*. Their contralaterally projecting axons suggest that these cells might play a role in coupling both brain hemispheres, like the PDF-*ir* neurons in the circadian clock of various insects^4–7,21,23^. We caution, however, that the involvement of PDFs in the function of the circadian clock of tardigrades remains to be demonstrated.

Localization of *pdfr*-expressing target cells in all major tissues and organs of *H. exemplaris*^46^ further suggests that PDFs might control or synchronize various functions in tardigrades, such as detection of light, neural processing, locomotion, feeding, digestion, osmoregulation, growth, development, oogenesis/reproduction, and formation of stylets and claws—a process related to molting^47^. The role of PDF/PDFR signaling in storage cells is unclear, but since these cells have been mainly associated with the production, storage and transport of lipids, polysaccharides and proteins^48,49^, PDFs might additionally regulate the storage and distribution of nutrients in the tardigrade body. This wide variety of functions parallels the reported multiple roles of PDFs in nematodes, ranging from the control of locomotion and mate searching to mechano- and chemosensation, including the sensation of oxygen^19,29,30^. A recent study further revealed that PDFR is expressed in multiple organs of the fruit fly *Drosophila melanogaster*^50^, suggesting hormonal control and multiple functions of the sole PDF, which like the three PDFs of *H. exemplaris* has originated from *pdf-I* of the ecdysozoan ancestor^12^. Similar observations, though without functional analyses, have been made in onychophorans^11^. These findings, together with our present observations, suggest a neuroendocrine role and multifunctionality of PDFs in the last common ancestors of Panarthropoda and Ecdysozoa. Focusing on multiple rather than just clock-related roles of PDFs in distantly related species would help to clarify the variety of functions and evolution of PDF/PDFR systems across protostomes.

## Methods

### Rearing and collection of specimens

Specimens of the tardigrade species *Hypsibius exemplaris* Ga[siorek *et al*., 2018^33^ (Eutardigrada, Hypsibiidae; strain Z151, Sciento, UK) were maintained under a 12/12 h light/dark cycle regime and fed with algae (*Chlorococcum* sp.) as described previously^51^. Approximately 50 adult specimens were collected with a glass pipette under a stereomicroscope for each localization experiment (immunolabeling and *in situ* hybridization), whereas several hundred specimens were extracted using a gauze (mesh size: 50 µm) for RNA extraction and purification.

### Transcriptomic, genomic, and phylogenetic analyses

The coding sequences of three *pdf* homologs (*He-pdf-1*: KP266565; *He-pdf-2*: KP266566; and *He-pdf-3*: KP266567) of *H. exemplaris* were obtained from the publicly available NCBI (National Center for Biotechnology Information) repository^12^. To clarify the orthology and evolutionary history of *pdf* genes across eutardigrades, we searched for putative *pdf* orthologs in the assembled transcriptomes from several tardigrade species using either tBLASTn or BLASTp v2.12.0+^52^ and the three *pdf* homologs of *H. exemplaris* as queries (Supplementary Table 1). We further applied a previous methodology^12^ to obtain the best maximum likelihood tree using RAxML v8.2.12^53^ and raxmlGUI v2.0.10^54^, but expanded the dataset by including *pdf* sequences from additional species of tardigrades and other panarthropods and aligned all sequences using the MAFFT online version^55^(G-INS-i option) (Supplementary Table 1). After inferring the best tree from 100 independent runs under empirical LG+G substitution model (according the model test implemented in raxmlGUI), bootstrap support values were estimated from 100 pseudoreplicates of the original alignment. The resulting tree was visualized using iTol v6^56^ and edited with Illustrator CS5.1 (Adobe Inc., San Jose, CA, USA).

Available transcriptome assemblies of *H. exemplaris* (GenBank accession numbers: GBZR01000000, GFGW01000000, and GJGU01000000) were used to search for putative *pdfr* transcripts. Initially, tBLASTn v2.12.0+ searches^52^ were conducted against all three assemblies using sequences from the onychophoran *Euperipatoides rowelli* (GenBank accession number: MT080366.1) and the fruit fly *Drosophila melanogaster* (NP_570007.2) as query sequences. Afterwards, the putative *pdfr* transcript of *H. exemplaris* was used as query for BLAST searches in genomic scaffolds of *Ramazzottius varieornatus* (BDGG00000000.1) and *Paramacrobiotus metropolitanus* (BHEN00000000.1) to obtain the putative *pdfr* genes of the respective species. For heterotardigrades, the transcriptome assemblies of *Echiniscus testudo* and *Echiniscoides sigismundi* were obtained from publicly available databases and an unpublished genome assembly of *Batillipes* sp. (Supplementary Table 1). Thereafter, tBLASTn v2.12.0+ searches^52^ were performed using the putative *pdfr* sequences from *H. exemplaris* and the previously reported *pdfr* sequence from *Euperipatoides rowelli*^11^ as bait sequences. The blast search yielded 21 hits for *Echiniscus testudo*, 20 hits for *Echiniscoides sigismundi* and eight hits for *Batillipes* sp. Subsequently, reciprocal BLAST searches with the resulting hits against the nucleotide database (nr/nt) of NCBI were performed to identify putative *pdfr* genes from heterotardigrades.

The transmembrane domains of putative *pdfr* sequences of *H. exemplaris*, *Paramacrobiotus metropolitanus*, *Echiniscus testudo*, *Echiniscoides sigismundi* and *Batillipes* sp. were predicted using the SMART online database^57^ and included in a previous dataset of ∼1,000 class B G protein-coupled receptors obtained from a cluster analysis of ∼18,000 bilaterian G protein-coupled receptors, including PDFRs^11^. After aligning the dataset using the MAFFT online version^55^ (G-INS-i option) a combined maximum likelihood analysis of 10 independent inferences and 100 thorough bootstrap pseudoreplicates was conducted under a dataset specific GTR+G model using RAxML v8.2.12^53^ and raxmlGUI v2.0.10^54^. The resulting tree was visualized using iTol v6^56^ and edited with Illustrator CS5.1 (Adobe Inc.). To analyze the genomic structure and to clarify whether there are any additional copies of the studied *pdf* and *pdfr* genes, BLASTn searches against the whole genomes of *H. exemplaris* (MTYJ00000000.1) and *Ramazzottius varieornatus* (BDGG00000000.1) were performed and intron/exons identified in the obtained genomic scaffolds by direct comparison of transcriptomic and genomic sequences of the regarding genes (Supplementary Data 1).

### Amplification and cloning of *pdf* and *pdfr* fragments

Total RNA was isolated from several hundred specimens of *H. exemplaris* with TRIzol^®^ reagent (Thermo Fisher Scientific) and resulting RNA was purified with RNeasy^®^ MinElute^®^Cleanup Kit (Qiagen, Hilden, Germany) according to manufacturer’s protocols. The concentration of the purified RNA was measured with a Qubit 3.0 Fluorometer (Life Technologies). The first-strand cDNA synthesis from purified RNA was performed using random hexamer primers (3 μg/mL) and SuperScript IV reverse transcriptase (Thermo Fisher Scientific, Waltham, MA, USA) according to the manufacturer’s instruction. Subsequently, full-length coding sequences of the target genes were amplified with gene-specific forward and reverse primers from previously obtained first strand cDNA (Supplementary Table 2). The amplicons of *He*-*pdf-1, He*-*pdf-2* and *He*-*pdf*-*3* were ligated into pJET1.2 vector (Thermo Fisher Scientific) as per manufacturer’s specification and transferred to competent *E. coli* TOP10 cells for overnight growth in lysogeny broth plates at 37 °C. Several colonies were picked to perform colony PCR and selected clones were verified by Sanger sequencing (Eurofins Genomics GmbH, Ebersberg, Germany). The amplification by designing gene-specific primers, cloning, and sequencing of the *He*-*pdfr* fragments were conducted using the same approach to obtain full-length coding sequences (Supplementary Table 2).

### Quantitative analysis of *pdf* and *pdfr* transcripts

The abundance of the identified *He-pdf-1*, *He-pdf-2*, *He-pdf-3*, and *He-pdfr-A/B* transcripts was estimated using segemehl 0.3.4^58^ (Bioinformatics, University of Leipzig, Germany) as described previously^59^. In brief, the raw sequence reads from four independent RNA-seq experiments of *H. exemplaris* (Sequence Read Archive run accession numbers: SRR14868527, SRR5218239, SRR5218240, and SRR5218241) were mapped back on the corresponding sequences allowing for a maximum of 5 % mismatch of nucleotides per read (Supplementary Data 2). Subsequently, the relative abundance (matched nucleotides per position and giga base pair) of genes was normalized using the relative abundance of the housekeeping gene *glyceraldehyde-3-phosphate dehydrogenase* (*GAPDH*) as a reference (100 %). We have chosen this gene, as it shows relatively low variation among the studied transcriptomes. The results were visualized as box plots at linear and logarithmic scales.

### Matrix Assisted Laser Desorption Ionization – Time of Flight Mass Spectrometry (MALDI-TOF MS)

Tardigrades were anesthetized on ice. Heads were opened and covered with ice-cold physiological insect saline (128 mM NaCl, 2.7 mM KCl, 2 mM CaCl_2_, 1.2 mM NaHCO_3_, pH 7.25). For direct tissue profiling experiments (n=15), brains were removed, subdivided in smaller portions using ultrafine scissors and transferred into a drop of water placed on a sample plate for MALDI-TOF MS analysis using a glass capillary fitted to a tube with a mouthpiece as described elsewhere^60^. Immediately after transfer, water was removed around the sample and the tissue was allowed to dry before matrix application.

For brain extraction (n=2), two samples consisting of 100 brains each were prepared. Brains were dissected as described above and transferred with a glass capillary fitted to a tube with a mouthpiece into 30 μL extraction solution on ice. Sample #1 contained 50 % methanol, 49 % H_2_O and 1 % formic acid (FA), whereas sample #2 contained 90 % ethanol, 9 % H_2_O and 1 % acetic acid. Tissue samples were homogenized using an ultrasonic bath (Transonic 660/H, Elma Schmidbauer GmbH, Hechingen, Germany) for 90 min on ice. Afterwards, the samples were centrifuged for 15 min at 13,000 rpm at 4 °C. The supernatants were separated and then evaporated in a vacuum concentrator to remove organic solvent. Extracts were stored at −20 °C until use. For matrix application, we followed a previous protocol^61^, except that we applied only 10 mg/mL 2,5–dihydroxybenzoic acid (DHB; Sigma- Aldrich, Steinheim, Germany) dissolved in 20 % acetonitrile, 1 % FA and 79 % H_2_O (Fluka) as matrix. For direct tissue profiling, dried tissue samples were covered with 0.1 µL 2,5- Dihydroxybenzoic acid solution applied with a 0.1–2.5 µL pipette (Eppendorf AB, Hamburg, Germany). For extract analysis, 0.1 µL concentrated supernatant was mixed with 0.1 µL DHB solution and again applied with a 0.1–2.5 µL Eppendorf pipette. Subsequently, sample spots were dried with a commercially available hair dryer to form homogeneous crystals.

Mass spectra were acquired manually using an ultrafleXtreme TOF/TOF mass spectrometer (Bruker Daltonik GmbH, Bremen, Germany) in reflector positive ion mode in a mass range of 900–3,000 Da. For calibration, the following mixture of synthetic peptides was used: (i) short neuropeptide F receptor of *Drosophila melanogaster*, (Drm)-sNPF-1[4–11]; (ii) periviscerokinin-1 of *Locusta migratoria* (Lom)-PVK-1; (iii) FMRFamide-12 of *Periplaneta americana*, (Pea)-FMRFa-12; (iv) allatotropin of *Manduca sexta* (Mas)-AT; (v) IPNamide of *Drosophila melanogaster*, (Drm)-IPNa; (vi) SKN of *Periplaneta americana*, Pea-SKN; and (vii) human glucagon. Laser fluency was adjusted to provide an optimal signal-to-noise ratio. The obtained data were processed using FlexAnalysis V.3.4 software package (Bruker Daltonik GmbH).

### Quadrupole Orbitrap MS Coupled to Nanoflow HPLC

For fragmentation analysis and sequence evaluation, brain extracts were desalted using self-packed Stage Tip C18 (IVA Analysentechnik e.K., Meerbusch, Germany) spin columns^62^ before injecting the samples into the nanoLC system. For analysis, peptides were separated on an EASY nanoLC 1000 UPLC system (Thermo Fisher Scientific, Waltham, MA) using 50 cm RPC18 columns (fused Silica tube with ID 50 μm ± 3 μm, OD 150 μm ± 6 μm, Reprosil 1.9 μm, pore diameter 60 Å, Dr. Maisch, Ammerbuch-Entringen, Germany) and a binary buffer system (A: 0.1 % FA, B: 80 % ACN, 0.1 % FA) as described for *Cataglyphis nodus* samples^61^. Running conditions were as follows: linear gradient from 2 % to 62 % B in 110 min, 62 % to 75 % B in 30 min, and final washing from 75 % to 95 % B in 6 min (45 °C, flow rate 250 nL/min). Finally, the gradients were re-equilibrated for 4 min at 5 % B. High Performance Liquid Chromatography (HPLC) was coupled to a Q-Exactive Plus (Thermo Scientific, Bremen, Germany) mass spectrometer. MS data were acquired in a top-10 data-dependent method dynamically choosing the most abundant peptide ions from the respective survey scans in a mass range of 300−3,000 m/z for higher-energy collisional dissociation (HCD) fragmentation. Full mass spectrometry acquisitions ran with a resolution of 70,000, automatic gain control target (AGC target) at 3e6, and maximum injection time at 80 ms. HCD spectra were measured with a resolution of 35,000, AGC target at 3e6, maximum injection time at 240 ms, 28 eV normalized collision energy, and dynamic exclusion set at 25s. The instrument was run in peptide recognition mode (i.e., from two to eight charges), singly charged and unassigned precursor ions were excluded. Raw data were analyzed with PEAKS Studio 10.5 (BSI, ON, Canada). Neuropeptides were searched against an internal database comprising neuropeptide precursor sequences from tardigrades with parent mass error tolerance of 0.2 Da and fragment mass error tolerance of 0.2 Da. Setting enzymes: none was selected because samples were not digested. The false discovery rate (FDR) was enabled by a decoy database search as implemented in PEAKS 10.5. Following posttranslational modifications (PTM) were selected: C-terminal amidation as fixed PTM and oxidation at methionine, phosphorylation, sulfation as variable PTMs. In each run a maximum of three variable PTMs per peptide were allowed. Fragment spectra with a peptide score (−10 lgP) equivalent to a P-value of ∼1 %, were manually reviewed.

### Bioluminescence Resonance Energy Transfer (BRET) assays

We tested the functionality and potential differences in interaction of the He-PDF-1, He- PDF-2 and He-PDF-3 peptides with both splice variants of their receptor, He-PDFR-A and He-PDFR-B. For this purpose, we established BRET assays using an Epac-based sensor (Epac-L) to examine cAMP responses in transfected human cells (HEK293T) as described previously^11^ for the onychophoran *Euperipatoides rowelli.* A POLARstar Omega microplate reader (BMG Labtech, Cary, NC, USA) was used to measure the resulting raw luminescence responses after stimulating the transfected cells with different concentrations (10^−1^^1^–10^−5^ M) of synthetic PDFs^11^ (Biomatik Corp., Kitchener, Ontario, Canada) (Supplementary Data 1). Their ratios [emission acceptor (515 nm)/emission donor (410 nm)]^63^ were box plotted against the respective PDF concentration (n=8 each) in dose response curves. The control experiments were conducted with cells expressing Epac-L alone (−*He-pdfr*-*A/B* in Fig. 2c, d), stimulation without PDFs (−PDF in Fig. 2c, d), and exposure to Forskolin (50 μM final concentration; Sigma-Aldrich Chemie GmbH, Munich, Germany) and IBMX (100 μM final concentration; Sigma-Aldrich Chemie GmbH), respectively (+Forsk./IBMX in Fig. 2c, d).

### Synthesis of antibodies and specificity tests

In a previous study^12^, specimens of *H. exemplaris* (formerly referred to as *H. dujardini*^33^) failed to stain with a polyclonal antiserum raised against the synthetic β-PDH of the crustacean *Uca pugilator* (code 3B3; Dircksen *et al.*^15^; catalog No. PDH beta; RRID: AB_231509), which has been widely used to localize PDHs/PDFs in a variety of animals^4,12,15,22^. We therefore generated customized polyclonal antibodies against each of the three mature PDFs of *H. exemplaris* as described previously for the onychophoran *Euperipatoides rowelli*^11^. The antibodies (anti-He-PDF-1: IG-P1037, LOT#2080B1, 21 µg/mL; anti-He-PDF-2: IG-P1038, LOT# 2025B2, 22 µg/mL; and anti-He-PDF-3: IG-P1039, LOT# 2082B, 31 µg/mL) (Supplementary Table 3) were commercially produced and purified (immunoGlobe GmbH, Himmelstadt, Germany) as described previously^11^. To clarify the spatial relationship between parts of the visual system (eye) and PDF-*ir* cells, we further generated a specific antibody against the visual rhabdomeric opsin of *H*. *exemplaris* (anti-He- R-Opsin-V: ID-Nr. 78-01-15; 0.8 mg/mL), based on the previously identified *r-opsin-v* sequence (formerly referred to as “*Hd-r-opsin Hypsibius dujardini*”^64^; GenBank accession number: KM086335). The antibody was commercially synthetized and purified by Peptide Specialty Laboratories GmbH, Heidelberg, Germany (Supplementary Table 3).

For testing the specificity of the customized antibodies as well as the previously used polyclonal antiserum against the β-PDH peptide of *Uca pugilator*, western blots were performed using synthetic He-PDF-1, He-PDF-2, and He-PDF-3 peptides (Biomatik Corp.). First, gradient acrylamide Tris/Tricine gels (4 %–10 %–16 %) were prepared^65^ and each well was loaded with 400 ng of the corresponding peptide, mixed with 2× Laemmli sample buffer (0.125 M Tris base, 0.14 M SDS, 20 % glycerol, 10 % 2-mercaptoethanol, Orange-G; pH 6.8). One well was loaded with 6 µL of a protein standard (Spectra^TM^ Multicolor Low Range Protein Ladder; Thermo Fisher Scientific). The peptides were separated in the gels by SDS-PAGE and subsequently transferred to PVDF membranes with 0.2 μm pore size (Carl Roth GmbH & Co. KG, Karlsruhe, Germany) using the semi-dry western blot technique. The membranes were blocked with 4 % bovine serum albumin (BSA; Carl Roth GmbH & Co. KG) in tris-buffered saline containing Tween 20 (TBS-T; 0.02 M Tris base, 0,15 M sodium chloride, 0.05 % Tween 20; pH 7.4) for 60 min and incubated with primary antibodies (anti-He-PDF-1: 1 µg/mL; anti-He-PDF-2: 1 µg/mL; anti-He-PDF-3: 1.5 µg/mL; diluted in TBS-T containing 1 % BSA) overnight at 4 °C. On the following day, after a few rinses with TBS-T, the membranes were incubated with peroxidase-conjugated goat anti-rabbit IgG (1:5,000; Jackson ImmunoResearch Laboratories Inc., Philadelphia, USA) at room temperature for three hours. Following multiple rinses with TBS-T, a chemiluminescent substrate (Pierce^TM^ ECL Western Blotting Substrate; Thermo Fisher Scientific) was added prior to detection. Finally, the Odyssey^®^ Fc Imager (LI-COR Biosciences, Nebraska, USA) was used for imaging the membranes. Brightness and contrast of acquired images were adjusted using Image Studio Version 6.0.0.28 (LI-COR Biosciences).

### Immunohistochemistry

The preparation of specimens for immunohistochemistry, including asphyxiation, puncturing and enzyme treatment, was performed as described previously^51^, except that we used a different fixative^66^ (4 % PFA, 7.5 % picric acid in 0.1 M PBS, pH 7.4) for fixation. After several rinses with PBS-Tx (PBS, 1 % Triton X-100), specimens were incubated in blocking solution containing 10 % normal goat serum (NGS; Sigma-Aldrich Chemie GmbH) in PBS- Tx for 1h at 4 °C. Thereafter, primary antibodies [either rabbit anti-He-PDF-1 (0.125 µg/mL in PBS-Tx, 1 % NGS), rabbit anti-He-PDF-2 (1 µg/mL in PBS-Tx, 1 % NGS) or rabbit anti-He-PDF-3 (3.1 µg/mL in PBS-Tx, 1 % NGS)] were applied and incubated for 3 days at 4 °C. Following multiple washes in PBS-Tx, a fluorescently conjugated secondary antibody [either goat anti-rabbit Alexa Fluor^®^ 488 (1:2000; Thermo Fisher Scientific) or goat anti-rabbit Alexa Fluor^®^ 568 (1:2000; Thermo Fisher Scientific)] were applied for 2 days at 4 °C (Supplementary Table 4). After several rinses with PBS-Tx, specimens were incubated in a solution containing the DNA marker DAPI (4′,6-diamidino-2-phenylindole; 1 ng/mL; Carl Roth GmbH & Co. KG) for 1 h at room temperature and then mounted in ProLong^TM^ Gold Antifade Mountant (Thermo Fisher Scientific) between two cover slips.

For double immunolabeling, two sets of experiments were conducted, combining either rabbit He-PDF-1 with rabbit He-PDF-2 or rabbit He-PDF-1 with rabbit He-PDF-3 antibodies, respectively. Beyond these modifications, we followed a previous protocol^11^ for double immunolabeling. Affinipure Fab Fragments (goat anti-rabbit; 40 μg/mL; Jackson ImmunoResearch Laboratories Inc., USA) in a solution of PBS-Tx with 1 % bovine serum albumin (BSA) were used for double immunolabeling to distinguish each two PDFs (Supplementary Table 5). The same concentrations of primary antibodies were used as for single immunolabeling and incubation proceeded for 2 days. We further performed double immunolabeling with the He-R-Opsin-V (10 µg/mL in PBS-Tx, 1 % NGS) and the He-PDF-1 (0.125 µg/mL in PBS-Tx, 1 % NGS) antibodies (Supplementary Table 3), using both simultaneously. The methodology was the same as for single immunolabeling.

### *In situ* hybridization

For mRNA detection using Hybridization Chain Reaction – Fluorescence *In Situ* Hybridization (HCR-FISH), split-initiator probes for *He-pdf-1*, *He-pdf-2*, *He-pdf-3, He-pdfr* and *He-r-opsin-v* were synthesized by Molecular Instruments, Los Angeles, USA (Supplementary Table 6). We followed a previous protocol^67^ for preparing the specimens of *H*. *exemplaris* for conducting HCR-FISH with the following alterations: Asphyxiated specimens were fixed in 4 % PFA (in PBS, pH 7.4) for 1.5 h. Fixed specimens were incubated with Proteinase K (Thermo Fisher Scientific; 0.25 mg/mL in PBS-Tw [PBS with 1% Tween 20]) for 30 min at room temperature. Thereafter, HCR-FISH was performed as described previously^68^ with the following modifications: Fixed specimens were first incubated with the probes for three days at 37 °C and then with the amplifier reagent conjugated to a fluorophore for two days at room temperature. The specimens were counterstained with DAPI for 1 h, rinsed several times in PBS and mounted between two cover slips in ProLong^TM^ Gold antifade Reagent (Thermo Fisher Scientific). Double HCR-FISH was conducted by simultaneously applying two probes. Double labeling of mRNA and the respective protein/peptide was performed by combining HCR-FISH with immunostaining. To achieve this, first protein detection and then mRNA detection were performed following a previous protocol^68^ with the following modification: The incubation with the initiator-labeled secondary antibodies was carried out overnight. Negative control experiments were performed by leaving out the probes but adding the respective amplifiers. Positive control experiments were conducted by using sets of redundant probes^69^ synthesized by Molecular Instruments.

In addition to HCR-FISH, we performed classical whole-mount *in situ* hybridization using digoxigenin-labeled RNA probes (*He*-*pdf-1*, *He*-*pdf-2* and *He*-*pdf-3*; 1–3 ng/μL each) according to a previous protocol^67^ with the same modifications described above for HCR-FISH. Hybridization was carried for three days at 60–63 °C. Anti-DIG AP Fab Fragments antibody (Roche, Mannheim, Germany; diluted 1:4,000 in 2 % NGS in PBS-Tw) was applied for three days at 4 °C. After several rinses with PBS-Tw, chromogenic staining was done either with NBT/BCIP (Thermo Fisher Scientific) solution or Fast Red Kit (Sigma-Aldrich, USA) in the dark and the reaction was stopped when desired intensity of signal was achieved. Specimens were mounted in Fluoromount-G^TM^ (Thermo Fisher Scientific) and imaged under a bright-field microscope.

### Microscopy, three-dimensional reconstructions, and panel design

Specimens were analyzed using a confocal laser scanning microscope with an Airyscan module (Zeiss LSM 880; Carl Zeiss Microscopy GmbH, Jena, Germany). Raw Airyscan datasets were processed with the ZEN 2 imaging software (black edition; Carl Zeiss Microscopy GmbH). Brightness, contrast, and sharpness of z-stack images were adjusted in Fiji v.1.52^70^. The LUT option was used in Fiji for assigning false colors to stained structures. Three-dimensional (3D) models were created based on selected high-resolution confocal datasets using a segmentation tool and volume rendering in Amira 6.0.1 (Thermo Fisher Scientific). Specimens labeled with the classical non-fluorescence chromogenic stain (NBT/BCIP and Fast Red Kit) were imaged under a bright-field microscope (Axio Imager.M2; Carl Zeiss Microscopy GmbH) equipped with a digital camera (Axiocam 503 color; Carl Zeiss Microscopy GmbH). Panels and diagrams were designed with Photoshop 24.5.0 and Illustrator 27.6.1 (Adobe Inc.).

## Supporting information

Dutta_et_al_supplementary material

## Data availability

The original sources of transcriptomic and genomic assemblies used in this study are listed in the Supplementary Materials (Supplementary Table 1). The raw data used for this article will be made available by the authors on request. Sequenced clones of *He-pdfr-A* and *He-pdfr-B* were deposited in GenBank under the accession numbers 0056 and 0057.

## Authors’ contributions

GM, LH, SN, MS and FWH designed the research. LH and SD accomplished transcriptomic, genomic, and phylogenetic analyses. KA provided unpublished genomic assembly of *Batillipes* sp. LH conducted BRET experiments. SD, NM performed western blotting. MMG and SD carried out immunolabeling. SN performed mass spectrometry. SD executed *in situ* hybridization and 3D reconstructions. SD, LH, MMG, SN and GM analyzed the data. SD, LH, NM, SN and GM wrote the first draft. All authors have read and approved the final manuscript.

## Acknowledgments

The authors are thankful to Achim Werckenthin for initial discussions, Shinta Fujimoto for collecting specimens of *Batillipes* sp., Alexander Meier for providing the Epac-L cAMP sensor, Sonja Fuhrmann, Thordis Arnold, Erik Machal, and Irmtraud Hammerl-Witzel for assistance with laboratory work, Christine Martin for help with confocal microscopy, Michael Klinger for support with 3D reconstructions, and all members of the Department of Zoology at the University of Kassel for support with animal husbandry.

## Supplementary figure legends

**Supplementary** Fig. 1 **Maximum likelihood tree under LG+G substitution model of 110 known ecdysozoan PDF/PDH peptides.** Nematode PDF paralogs are boxed in light and dark blue, two onychophoran PDF paralogs in red and orange (PDF-I and PDF-II), and tardigrade PDFs in yellow. Bootstrap support values greater than 50 % are denoted. Scale bar indicates substitutions per site.

**Supplementary Fig. 2** Maximum likelihood tree under dataset-specific GTR+G model of amino acid sequence evolution of 996 class B G protein-coupled receptors. Note position of pigment-dispersing factor receptors (PDFRs) of tardigrades inside PDFR clade of protostome PDFRs (red box). PDFR splice variants of *H. exemplaris*, He-PDFR-A and He- PDFR-B, are highlighted in red. Bootstrap support values greater than 50 % are denoted. Scale bar indicates substitutions per site.

**Supplementary** Fig. 3 **Specificity of antibodies used to detect PDFs of *H. exemplaris*.** Western blots with synthetic peptides using protein ladder (leftmost track of each membrane) as reference. (**a–c**) Note that specific He-PDF-1, He-PDF-2 and He-PDF-3 antibodies exclusively recognize respective peptides, as illustrated by single bands in each membrane. Asterisks indicate absence of signal in controls. (**d**) In contrast to specific antibodies, β-PDH serum against crustacean peptide does not react with any of the three PDFs of *H. exemplaris*.

**Supplementary** Fig. 4 **Details of varicose projections of lateral cells in the brain of *H. exemplaris*.** Double immunostaining of head region showing localization of He-PDF-1 and visual opsin (He-R-Opsin-V). Projections of confocal substacks in dorsal view. Anterior is up in both images. Immunoreactivities of He-PDF-1 and He-R-Opsin-V are illustrated in yellow and blue, respectively, DNA staining in gray. (**a**) Overview. Note that peripheral axons (arrows) emerge from pseudounipolar lateral cells to terminate near dorsal surface of rhabdomeric receptor of eye. (**b**) Detail of left region of image in **a**. Abbreviations: ILC inner lobe PDF-immunoreactive cells, LAC lateral PDF-immunoreactive cells, RHC rhabdomeric cell. Scale bars: 5 µm (**a, b**).

**Supplementary** Fig. 5 **Co-localization of PDFs in the brain of *H. exemplaris*.** Double immunostaining with He-PDF-1 and He-PDF-2 (**a–c**), and He-PDF-1 and He-PDF-3 antibodies (**d–f**). Maximum projections of confocal micrographs in dorsal view (**a–f**). Anterior is up in all images. DNA staining is illustrated in blue. Arrows point to each right pair of inner lobe cells, details of which are shown in insets (upper right corner of each image). Note that all three PDFs are co-localized in inner lobe cells and their varicosities (arrowheads). Abbreviation: ILC inner lobe PDF-immunoreactive cells. Scale bars: 5 µm (**a– f**) including insets.

**Supplementary** Fig. 6 Expression of PDF-1 peptide and *pdf-1* transcripts in the brain of ***H. exemplaris* at different Zeitgeber times.** Immunostaining (**a–c**) and HCR-FISH (**d–f**) of specimens collected during subjective day (ZT4, ZT8) and night (ZT16). Projections of confocal substacks of head region in dorsal view. Anterior is up in all images. Expression of He-PDF-1 peptide is represented in green and *He-pdf-1* mRNA in magenta. DNA labeling is illustrated in gray. Note that there are no noticeable differences in intensity of signal between time points. Abbreviations: ILC inner lobe PDF-immunoreactive cells, LAC lateral PDF-immunoreactive cells, MEC median PDF-immunoreactive cells. Scale bars: 5 µm (**a–f**).

**Supplementary** Fig. 7 **Localization of *pdf-1*, *pdf-2* and *pdf-3* mRNA in the brain of *H. exemplaris*.** Results of classical *in situ* hybridization using digoxigenin (DIG)-labeled RNA probes. Light micrographs. Anterior is up in all images. (**a–c**) Expression of *He-pdf-1*, *He- pdf-2* and *He-pdf-3* mRNA in inner lobe cells of the brain. Note that *He-pdf-1* and *He-pdf-2* were stained with NBT/ BCIP (blue in **a**, **b**), whereas *He-pdf-3* was stained overnight with Fast Red (red in **c**). Note also that this method fails to detect weak expression of *He-pdf-1* in lateral, median and ventral extracerebral cells. Abbreviations: ILC inner lobe PDF-immunoreactive cells, PHA pharynx. Scale bars: 10 µm (**a–c**).

**Supplementary** Fig. 8 **Co-localization of *pdf* transcripts in the brain of *H. exemplaris*.** Double mRNA labeling. Anterior is up in all images. DNA staining is represented in gray. (**a–f**) Projections of confocal substacks of heads in dorsal view. Expression of *He-pdf-1* is illustrated in magenta, *He-pdf-2* in blue and *He-pdf-3* in green. Note co-expression of all three *pdf* genes in inner lobe cells. Abbreviation: ILC inner lobe PDF-immunoreactive cells. Scale bars: 5 µm (**a–f**).

**Supplementary** Fig. 9 **Co-localization of He-PDF-1 peptide and *He-pdf-1* mRNA in the brain of *H. exemplaris*.** mRNA labeling combined with immunolabeling. Confocal micrographs in dorsal (**a–c**) and lateral views (**d–f**). Anterior is up in **a–c** and left in **d–f**. Expression of He-PDF-1 peptide is illustrated in magenta, *He-pdf-1* mRNA in green. DNA staining is represented in gray. Note co-localization of peptide (He-PDF-1) and mRNA (*He- pdf-1*) in the same set of cells, including somata of inner lobe cells and lateral cells. Abbreviations: ILC inner lobe PDF-immunoreactive cells, LAC lateral PDF-immunoreactive cells, MEC median PDF-immunoreactive cells, PLA pharyngeal placoids (visualized by autofluorescence). Scale bars: 5 µm (**a–c**) and 10 µm (**d–f**).

**Supplementary** Fig. 10 **Quadrupole orbitrap mass spectra generated from brain extracts of *H. exemplaris.*** Transcriptome-predicted *pdf* precursor sequences were used as database for evaluating amino acid sequences of PDFs using PEAKS 10.5 software package.

**Supplementary** Fig. 11 **Comparison of expression levels of two isoforms of *He-pdfr* gene of *H. exemplaris.*** Box-plotted logarithmic chart showing relative expression levels of both splice variants, *He*-*pdfr*-*A* and *He*-*pdfr*-*B*, compared to *He-pdf-1*, *He-pdf-2*, and *He-pdf-3*. Diagram is based on comparative analysis of four transcriptomes using short read mapping in segemehl 0.3.4, with housekeeping gene *GAPDH* as reference. Boxes correspond to interquartile ranges. Circle inside each box represents arithmetic mean.

**Supplementary** Fig. 12 **Localization of *He-pdfr* mRNA in ventral trunk ganglia and somatic muscles of *H. exemplaris*.** Labeling was performed using *He-pdfr* probe. Projections of confocal substacks of specimens in ventral view. Anterior is up in all images. Gray color illustrates DNA staining in **a–c** and autofluorescence of somatic muscle in **d**, which was achieved by considerable increase of brightness and contrast of UV channel. (**a– c)** Localization of *He-pdfr* in trunk ganglia 1, 2 and 4. (**d**) Localization of *He-pdfr* in ventral somatic muscle cell (arrow). Scale bars: 5 µm (**a–d**).

**Supplementary** Fig. 13 **Localization of *He-pdfr* and *He*-*r*-*opsin*-*v* in the eye of *H. exemplaris*.** Double mRNA labeling with *He-pdfr* (yellow) and *He-r-opsin-v* probes (blue). Maximum confocal projection of specimen in dorsal view. Anterior is up. DNA staining is represented in gray. Note co-localization of *He-pdfr* and *He-r-opsin-v* in rhabdomeric cell of each eye (arrows). Abbreviation: PHA pharynx, PLA pharyngeal placoids (visualized by autofluorescence). Scale bar: 10 µm.

**Supplementary** Fig. 14 **Localization of *He-pdfr* and *He-pdf-1* in the brain of *H. exemplaris*.** Double mRNA labeling with *He-pdfr* (yellow) and *He-pdf-1* probes (magenta). Projections of confocal substacks of specimens in dorsal view. Anterior is up in all images. DNA staining is represented in gray. (**a**) Note co-localization of *He-pdfr* and *He-pdf-1* in inner lobe cells, lateral cells, and median cells. (**b**) Detail of *He-pdf-1* inner lobe cells and lateral cell from left brain hemisphere of same specimen as in **a**. (**c**) Detail of another specimen showing co-localization of *He-pdfr* and *He-pdf-1* in median cells and inner lobe cells. Abbreviations: ILC inner lobe PDF-immunoreactive cells, LAC lateral PDF-immunoreactive cells, MEC median PDF-immunoreactive cells, PLA pharyngeal placoids (visualized by autofluorescence). Scale bars: 10 µm (**a**), and 5 µm (**b**, **c**).

**Supplementary** Fig. 15 **Positive control of *He-pdfr* mRNA expression in *H. exemplaris.*** Validation experiment was performed using two sets of redundant *He-pdfr* probes. Set A is illustrated in magenta, set B in yellow. DNA staining is represented in gray. (**a–c**) Projections of substacks of confocal micrographs. Anterior is up in all images. Note overlapping expression of both sets of probes. Abbreviation: BR brain. Scale bars: 5 µm (**a– c**).

## Supplementary data

Supplementary Data 1. Summary statistics and numerical data of *pdf* and *pdfr* genes in *Hypsibius exemplaris* as well as *Ramazzottius varieornatus* and raw luminescence response data obtained from BRET assays.

Supplementary Data 2. Raw data and calculations of the quantitative analysis of *pdf* and *pdfr* transcripts in *Hypsibius exemplaris*.

## Supplementary movies

Supplementary Movie 1. Localization of He-PDF-1 in *H. exemplaris* with particular focus on position of PDF-*ir* neurons in the brain.

Supplementary Movie 2. Localization of He-PDF-1 in *H. exemplaris* with particular focus on position of ventral extracerebral PDF-*ir* cells.

Supplementary Movie 3. Localization of *He-pdf-1* mRNA in *H. exemplaris* with particular focus on the brain.

Supplementary Movie 4. Localization of *He-pdf-1* mRNA in *H. exemplaris* with particular focus on ventral extracerebral cells.

