## Supplementary material for "Pigment-dispersing factor neuropeptides act as multifunctional hormones and modulators in tardigrades": Dutta_et_al_supplementary material: suppl_fig_10.pdf

**PDF-1**; *m/z* 1992.19; NAEVLNSLIGLPRLLKDK-$NH_2$

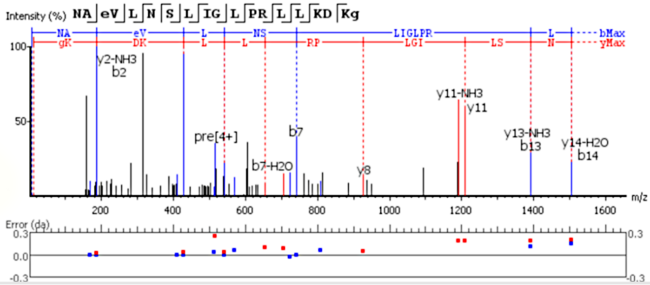


| # | b | b-H2O | b-NH3 | b (2+) | Seq | y | y-H2O | y-NH3 | y (2+) | # |
|---|---|---|---|---|---|---|---|---|---|---|
| 1 | 115.05 | 97.04 | 98.02 | 58.03 | N | | | | | 19 |
| 2 | 186.09 | 168.08 | 169.06 | 93.54 | A | 1949.19 | 1931.18 | 1932.16 | 975.10 | 18 |
| 3 | 329.15 | 311.14 | 312.12 | 165.07 | E(+14.02) | 1878.15 | 1860.14 | 1861.13 | 939.58 | 17 |
| 4 | 428.21 | 410.20 | 411.19 | 214.61 | V | 1735.10 | 1717.08 | 1718.07 | 868.05 | 16 |
| 5 | 541.30 | 523.29 | 524.27 | 271.15 | L | 1636.03 | 1618.02 | 1619.00 | 818.51 | 15 |
| 6 | 655.34 | 637.33 | 638.31 | 328.17 | N | 1522.94 | 1504.72 | 1505.92 | 761.97 | 14 |
| 7 | 742.37 | 724.40 | 725.35 | 371.69 | S | 1408.90 | 1390.89 | 1391.68 | 704.85 | 13 |
| 8 | 855.46 | 837.45 | 838.43 | 428.21 | L | 1321.87 | 1303.86 | 1304.84 | 661.43 | 12 |
| 9 | 968.54 | 950.53 | 951.51 | 484.77 | I | 1208.59 | 1190.77 | 1191.57 | 604.89 | 11 |
| 10 | 1025.56 | 1007.55 | 1008.54 | 513.24 | G | 1095.70 | 1077.69 | 1078.67 | 548.35 | 10 |
| 11 | 1138.65 | 1120.64 | 1121.62 | 569.75 | L | 1038.68 | 1020.67 | 1021.65 | 519.84 | 9 |
| 12 | 1235.70 | 1217.69 | 1218.67 | 618.35 | P | 925.54 | 907.58 | 908.57 | 463.30 | 8 |
| 13 | 1391.68 | 1373.79 | 1374.77 | 696.40 | R | 828.54 | 810.53 | 811.51 | 414.77 | 7 |
| 14 | 1504.72 | 1486.87 | 1487.86 | 752.94 | L | 672.44 | 654.32 | 655.41 | 336.72 | 6 |
| 15 | 1617.97 | 1599.96 | 1600.94 | 809.42 | L | 559.36 | 541.30 | 542.33 | 280.18 | 5 |
| 16 | 1746.06 | 1728.05 | 1729.04 | 873.53 | K | 446.27 | 428.21 | 429.25 | 223.64 | 4 |
| 17 | 1861.09 | 1843.08 | 1844.06 | 931.05 | D | 318.18 | 300.17 | 301.15 | 159.59 | 3 |
| 18 | 1989.19 | 1971.18 | 1972.16 | 995.09 | K | 203.15 | 185.14 | 186.09 | 102.08 | 2 |
| 19 | | | | | G(-.98) | 75.06 | 57.04 | 58.03 | 38.03 | 1 |

**PDF-2**; *m/z* 2078.21; NSEILNT**II**GLPNKLRQR-$NH_2$

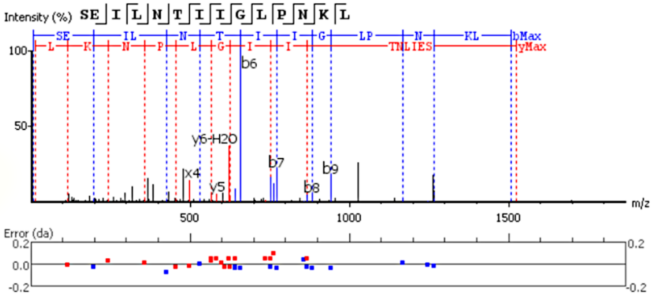


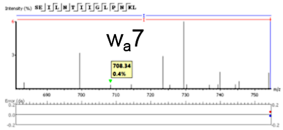


| # | b | b-H2O | b-NH3 | b (2+) | Seq | y | y-H2O | y-NH3 | y (2+) | # |
|---|---|---|---|---|---|---|---|---|---|---|
| 1 | 88.04 | 70.03 | 71.01 | 44.52 | S | | | | | 14 |
| 2 | 217.08 | 199.14 | 200.06 | 109.04 | E | 1437.87 | 1419.86 | 1420.84 | 719.43 | 13 |
| 3 | 330.17 | 312.16 | 313.22 | 165.58 | I | 1308.82 | 1290.81 | 1291.80 | 654.91 | 12 |
| 4 | 443.25 | 425.24 | 426.22 | 222.13 | L | 1195.74 | 1177.73 | 1178.71 | 598.36 | 11 |
| 5 | 557.29 | 539.28 | 540.27 | 279.15 | N | 1082.66 | 1064.65 | 1065.63 | 541.83 | 10 |
| 6 | 658.39 | 640.33 | 641.35 | 329.67 | T | 968.61 | 950.60 | 951.59 | 484.81 | 9 |
| 7 | 771.47 | 753.45 | 754.43 | 386.22 | L | 867.51 | 849.56 | 850.54 | 434.28 | 8 |
| 8 | 884.55 | 866.50 | 867.51 | 442.75 | I | 754.43 | 736.42 | 737.46 | 377.74 | 7 |
| 9 | 941.58 | 923.52 | 924.50 | 471.27 | G | 641.35 | 623.34 | 624.40 | 321.20 | 6 |
| 10 | 1054.61 | 1036.60 | 1037.59 | 527.81 | L | 584.33 | 566.32 | 567.35 | 292.69 | 5 |
| 11 | 1151.67 | 1133.66 | 1134.64 | 576.33 | P | 471.29 | 453.28 | 454.30 | 236.15 | 4 |
| 12 | 1265.72 | 1247.70 | 1248.68 | 633.37 | N | 374.24 | 356.22 | 357.21 | 187.62 | 3 |
| 13 | 1393.81 | 1375.79 | 1376.78 | 697.40 | K | 260.20 | 242.15 | 243.17 | 130.60 | 2 |
| 14 | | | | | L | 132.10 | 114.09 | 115.07 | 66.55 | 1 |

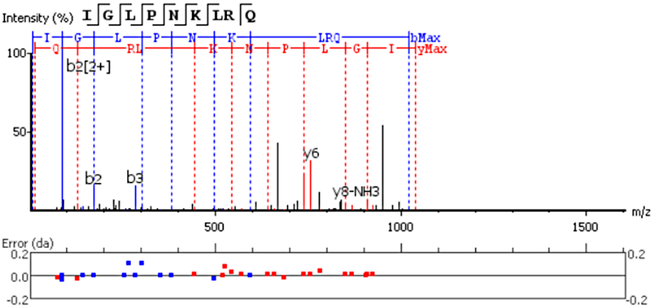


| # | b | b-H2O | b-NH3 | b (2+) | Seq | y | y-H2O | y-NH3 | y (2+) | # |
|---|---|---|---|---|---|---|---|---|---|---|
| 1 | 114.09 | 96.08 | 97.06 | 57.55 | I | | | | | 9 |
| 2 | 171.11 | 153.10 | 154.09 | 86.10 | G | 925.55 | 907.54 | 908.52 | 463.28 | 8 |
| 3 | 284.20 | 266.08 | 267.17 | 142.60 | L | 868.52 | 850.53 | 851.50 | 434.77 | 7 |
| 4 | 381.25 | 363.24 | 364.22 | 191.13 | P | 755.44 | 737.43 | 738.41 | 378.23 | 6 |
| 5 | 495.33 | 477.28 | 478.27 | 248.15 | N | 658.39 | 640.39 | 641.36 | 329.70 | 5 |
| 6 | 623.39 | 605.38 | 606.36 | 312.19 | K | 544.32 | 526.26 | 527.33 | 272.68 | 4 |
| 7 | 736.47 | 718.46 | 719.45 | 368.74 | L | 416.26 | 398.25 | 399.23 | 208.63 | 3 |
| 8 | 892.57 | 874.56 | 875.55 | 446.79 | R | 303.18 | 285.17 | 286.15 | 152.09 | 2 |
| 9 | | | | | Q | 147.08 | 129.10 | 130.05 | 74.06 | 1 |

**PDF-3**; *m/z* 2050.21; NSEILNT**LI**GLPNKLKQR-$NH_2$

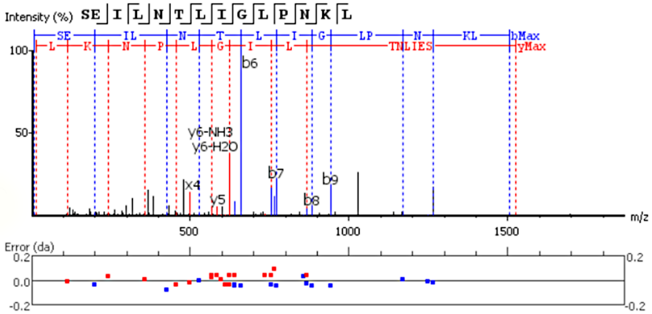


| # | b | b-H2O | b-NH3 | a | b (2+) | Seq | x | y | y-H2O | y-NH3 | y (2+) | # |
|---|---|---|---|---|---|---|---|---|---|---|---|---|
| 1 | 88.04 | 70.03 | 71.01 | 60.04 | 44.52 | S | | | | | | 14 |
| 2 | 217.08 | 199.11 | 200.06 | 189.09 | 109.04 | E | 1463.85 | 1437.87 | 1419.86 | 1420.84 | 719.43 | 13 |
| 3 | 330.17 | 312.16 | 313.14 | 302.17 | 165.58 | I | 1334.80 | 1308.82 | 1290.81 | 1291.80 | 654.91 | 12 |
| 4 | 443.25 | 425.24 | 426.31 | 415.26 | 222.13 | L | 1221.72 | 1195.74 | 1177.73 | 1178.71 | 598.35 | 11 |
| 5 | 557.29 | 539.28 | 540.27 | 529.30 | 279.15 | N | 1108.64 | 1082.66 | 1064.65 | 1065.63 | 541.83 | 10 |
| 6 | 658.39 | 640.38 | 641.35 | 630.35 | 329.67 | T | 994.59 | 968.61 | 950.60 | 951.59 | 484.81 | 9 |
| 7 | 771.47 | 753.41 | 754.43 | 743.43 | 386.21 | L | 893.55 | 867.52 | 849.56 | 850.54 | 434.28 | 8 |
| 8 | 884.55 | 866.50 | 867.52 | 856.48 | 442.75 | I | 780.46 | 754.43 | 736.42 | 737.46 | 377.74 | 7 |
| 9 | 941.58 | 923.52 | 924.50 | 913.54 | 471.27 | G | 667.38 | 641.35 | 623.34 | 624.41 | 321.20 | 6 |
| 10 | 1054.61 | 1036.60 | 1037.59 | 1026.62 | 527.81 | L | 610.39 | 584.33 | 566.32 | 567.32 | 292.69 | 5 |
| 11 | 1151.67 | 1133.66 | 1134.64 | 1123.67 | 576.33 | P | 497.30 | 471.29 | 453.28 | 454.30 | 236.15 | 4 |
| 12 | 1265.74 | 1247.72 | 1248.68 | 1237.72 | 633.36 | N | 400.22 | 374.24 | 356.22 | 357.21 | 187.62 | 3 |
| 13 | 1393.81 | 1375.79 | 1376.78 | 1365.81 | 697.40 | K | 286.18 | 260.20 | 242.15 | 243.13 | 130.60 | 2 |
| 14 | | | | | | L | 158.08 | 132.10 | 114.09 | 115.09 | 66.55 | 1 |

supplementary Fig. 10
