## Supplementary material for "Pigment-dispersing factor neuropeptides act as multifunctional hormones and modulators in tardigrades": Dutta_et_al_supplementary material: suppl_tables.docx

**Table 1: Data availability of eutardigardes and heterotardigrades, used in this study**

| **Taxon** | | **Species** | **Assembly** | **Data availability** |
| --- | --- | --- | --- | --- |
| Eutardigrada | Parachela | *Hypsibius exemplaris* | Transcriptome | GenBank accessions:  GBZR00000000.1  GFGW00000000.1  GJGU00000000.1 |
|  |  |  | Genome | MTYJ00000000.1 |
|  |  | *Ramazzottius varieornatus* | Genome | GenBank accession:  BDGG00000000.1 |
|  |  | *Richtersius coronifer*^1^ | Transcriptome | https://doi.org/10.6084/m9.figshare.8797184.v1 |
|  |  | *Paramacrobiotus richtersi* | Transcriptome | NCBI TSA:  GFGY00000000.1 |
|  |  | *Paramacrobiotus metropolitanus* | Genome | GenBank accession:  BHEN00000000.1 |
|  |  | *Mesobiotus philippinicus* | Transcriptome | https://doi.org/10.7910/DVN/CFNUGF/VCVLAO, Harvard Dataverse, V1 |
|  | Apochela | *Milnesium tardigradum* | Transcriptome | NCBI TSA:  GFGZ00000000.1 |
| Heterotardigrada | Echiniscoidea | *Echiniscus testudo* | Transcriptome | https://doi.org/10.7910/DVN/CFNUGF/6QPHBB, Harvard Dataverse, V1 |
|  |  |  | Genome | https://figshare.com/articles/dataset/Assembled_Genome_and_Transcriptome_of_heterotardigrade_Echiniscus_testudo/13060634 |
|  |  | *Echiniscoides* cf. *sigismundi* | Transcriptome | https://doi.org/10.7910/DVN/CFNUGF/NS9KRC, Harvard Dataverse, V1 |

1	Stec, D., Krzywański, Ł., Arakawa, K. & Michalczyk, Ł. A new redescription of *Richtersius coronifer*, supported by transcriptome, provides resources for describing concealed species diversity within the monotypic genus Richtersius (Eutardigrada). *Zoological Lett.* **6**, 2, (2020).

**Table 2: Specific primers used for amplification of gene fragments in this study**

| **Name of the genes**  **& accession no.** | **Forward and reverse primers** | **Sequence length (bp)** | **Annealing temperature (Tm)** |
| --- | --- | --- | --- |
| *He-pdf-1*  GenBank: KP266565.1 | For: 5ʹATGCAGTGTGTTACCCTCGC 3ʹ  Rev: 5ʹ TTATCGGCCCTTGTCCTTGA 3ʹ | 255 | 63°C |
| *He-pdf-2*  GenBank: KP266566.1 | For: 5ʹATGGATGTCAAAGCCTTTCTGTT 3ʹ  Rev: 5ʹ TTAGCCCCTTTGACGTAGTTTATTA 3ʹ | 294 | 63°C |
| *He-pdf-3*  GenBank: KP266567.1 | For: 5ʹ ATGGATTCACGCGTTCTTTTTTTTAC3ʹ  Rev: 5ʹ TCAACCTCGCTGTTTCAGCTTATTAG 3ʹ | 339 | 64°C |
| *He-pdfr-A/B*  GenBank: PQ050056.1 & PQ050057.1 | For: 5ʹ ATGATGCCGCCACCG 3ʹ  Rev: 5ʹ TCACATTTCAATTTCCTCCTTT 3ʹ | 1581 | 56°C |

**Table 3: List of customized primary antibodies which were used in this study with their peptide sequences, host species and manufacturer**

| **Name of the antibody** | **Peptide sequences, which were immunized** | **Host** | **Manufacturer** |
| --- | --- | --- | --- |
| Anti-He-PDF-1 | NAEVLNSLIGLPRLLKDK-NH_2_ | Rabbit | immunoGlobe GmbH, Himmelstadt, Deutschland  Product ID and Conc.  IG-P1037(25 µg/ml) |
| Anti-He-PDF-2 | NSEILNTIIGLPNKLRQR-NH_2_ | Rabbit | immunoGlobe GmbH, Himmelstadt, Deutschland  Product ID and Conc.  IG-P1038 (22 µg/ml) |
| Anti-He-PDF-3 | NSEILNTLIGLPNKLKQR-NH_2_ | Rabbit | immunoGlobe GmbH, Himmelstadt, Deutschland  Product ID and Conc.  IG-P1039 (31 µg/ml) |
| Anti-R-Opsin-Visual | CIKSPDEEIKGTSSVGKSTRIATGNGSS | Guinea pig | Peptide Specialty Laboratories GmbH, Heidelberg, Germany  ID-Nr. 78-01-15 (0.8 mg/ml) |

**Table 4: List of secondary antibodies, which were used in this study with their host species, concentration and manufacturers**

| **Name of the antibody** | **Concentration used for each experiment** | **Host** | **Manufacturer** |
| --- | --- | --- | --- |
| Anti-Rabbit-Alexa Fluor^®^-488 | 1: 2000 | Goat | Thermo Fisher Scientific, Waltham, USA |
| Anti-Rabbit-Alexa Fluor^®^-568 | 1: 2000 | Goat | Thermo Fisher Scientific, Waltham, USA |
| Anti-Guinea Pig-Alexa Fluor^®^-488 | 1:500 | Donkey | Thermo Fisher Scientific, Waltham, USA |
| Peroxidase-conjugated anti-Rabbit IgG | 1: 5000 | Goat | Jackson ImmunResearch Laboratories, Inc., USA |

**Table 5: List of commercially purchased primary antibodies which were used in this study with host species and manufacturers**

| **Name of the antibody** | **Concentration used for each experiment** | **Host** | **Manufacturer** |
| --- | --- | --- | --- |
| Anti-Rabbit-Fab-Fragments | 40 μg/mL | Goat | Jackson ImmunResearch  Laboratories, Inc., Pennsylvania, USA |
| Anti-Digoxigenin-AP, Fab fragments | 1: 2000 | Sheep | Roche |

**Table 6: List of probe and amplifiers for conducting HCR-FISH**

| **Target gene** | **Accession No.** | **Length**  **(nt)** | **Probe set size** | **Amplifiers** | **Probe working concentration** | **company** |
| --- | --- | --- | --- | --- | --- | --- |
| *He-pdf-1* | KP266565.1 | 255 | 4 | B1 | 4 µl to 500 µl hybridization buffer | Molecular Instruments, USA |
| *He-pdf-2* | KP266566.1 | 294 | 5 | B4 | 10 µl to 500 µl hybridization buffer |  |
| *He-pdf-3* | KP266567.1 | 339 | 5 | B3 | 13 µl to 500 µl hybridization buffer |  |
| *He-pdfr-11K10* | PQ050056.1 | 1581 | 20 | B3 | 15 µl to 500 µl hybridization buffer |  |
| *He-pdfr-11K10_*redundant set A | -- | -- | 10 | B2 | 10 µl to 500 µl hybridization buffer |  |
| *He-pdfr-11K10_* redundant set B | -- | -- | 10 | B5 | 10 µl to 500 µl hybridization buffer |  |
