## Supplementary figures and images for "Pigment-dispersing factor neuropeptides act as multifunctional hormones and modulators in tardigrades"

### suppl_fig_1.pdf

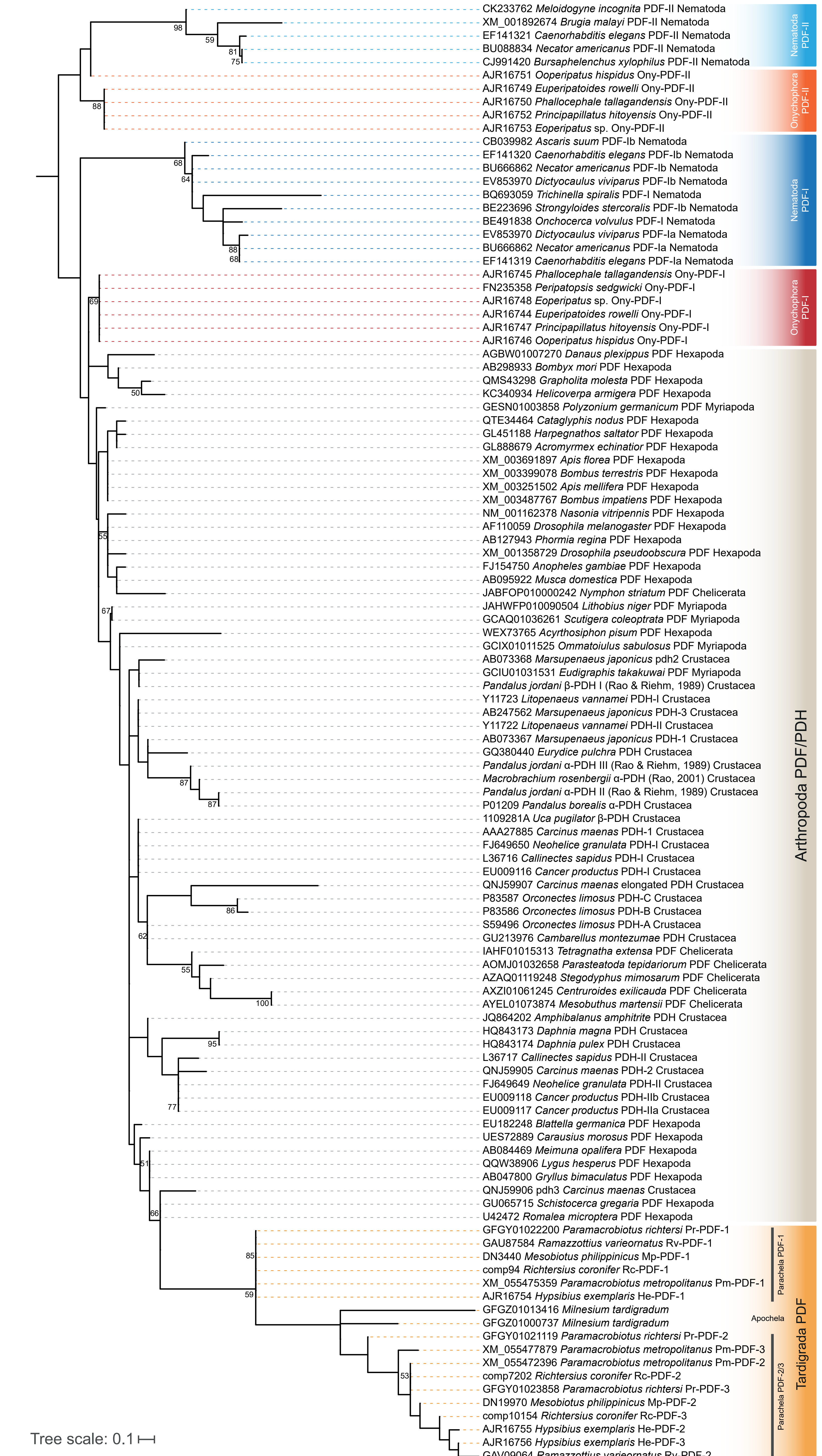


supplementary Fig. 1

### suppl_fig_2.pdf

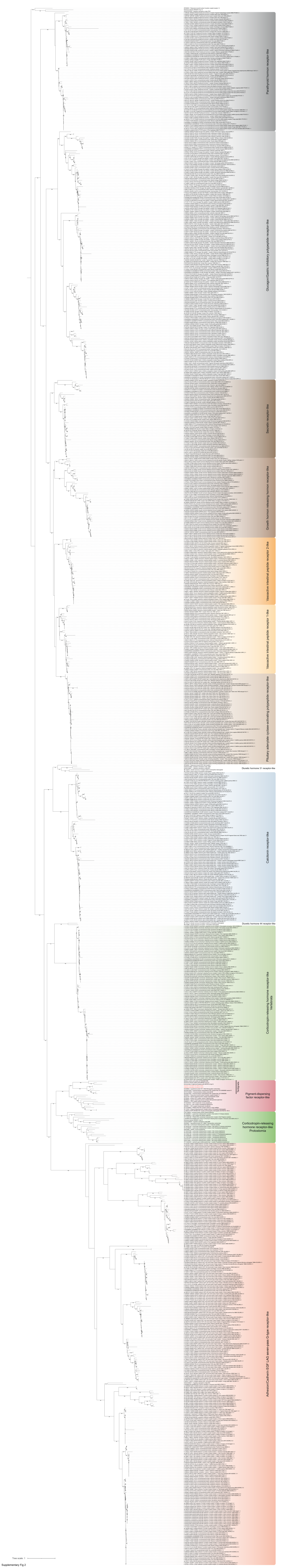


Supplementary Fig.2

### suppl_fig_3.pdf

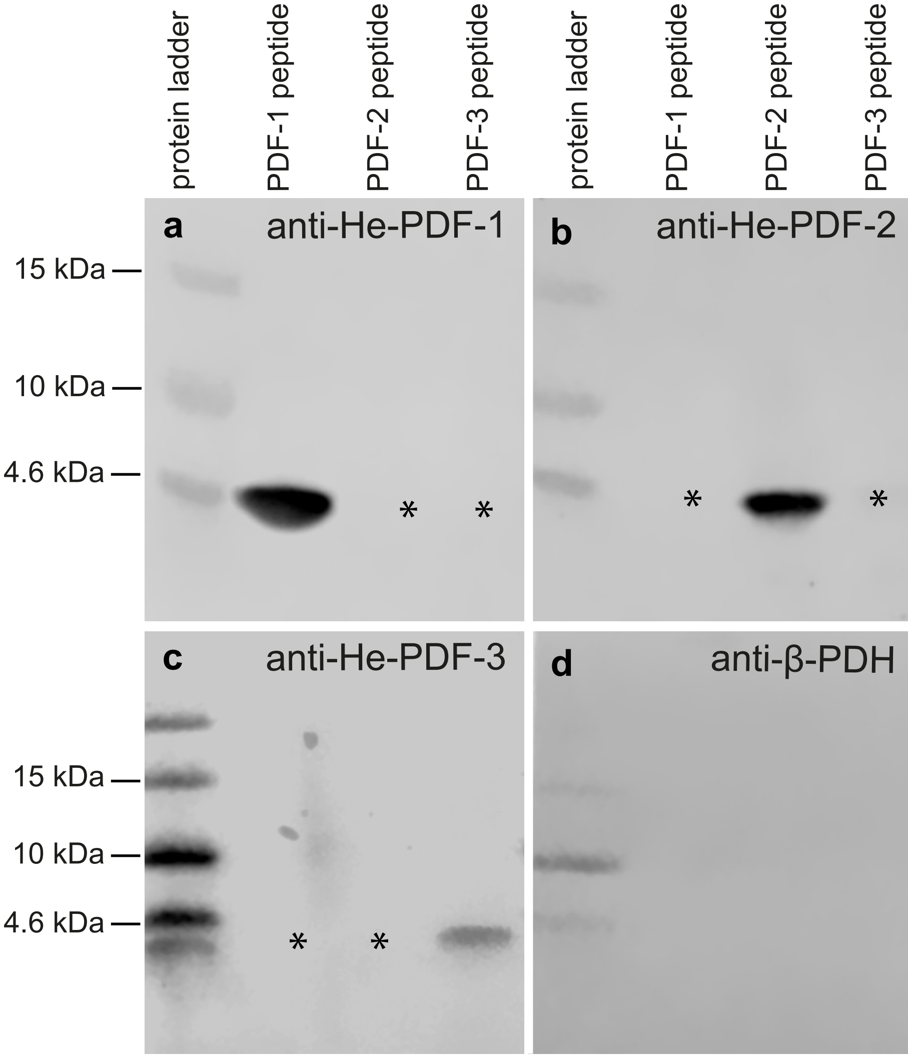


Supplementary Fig. 3

### suppl_fig_4.pdf

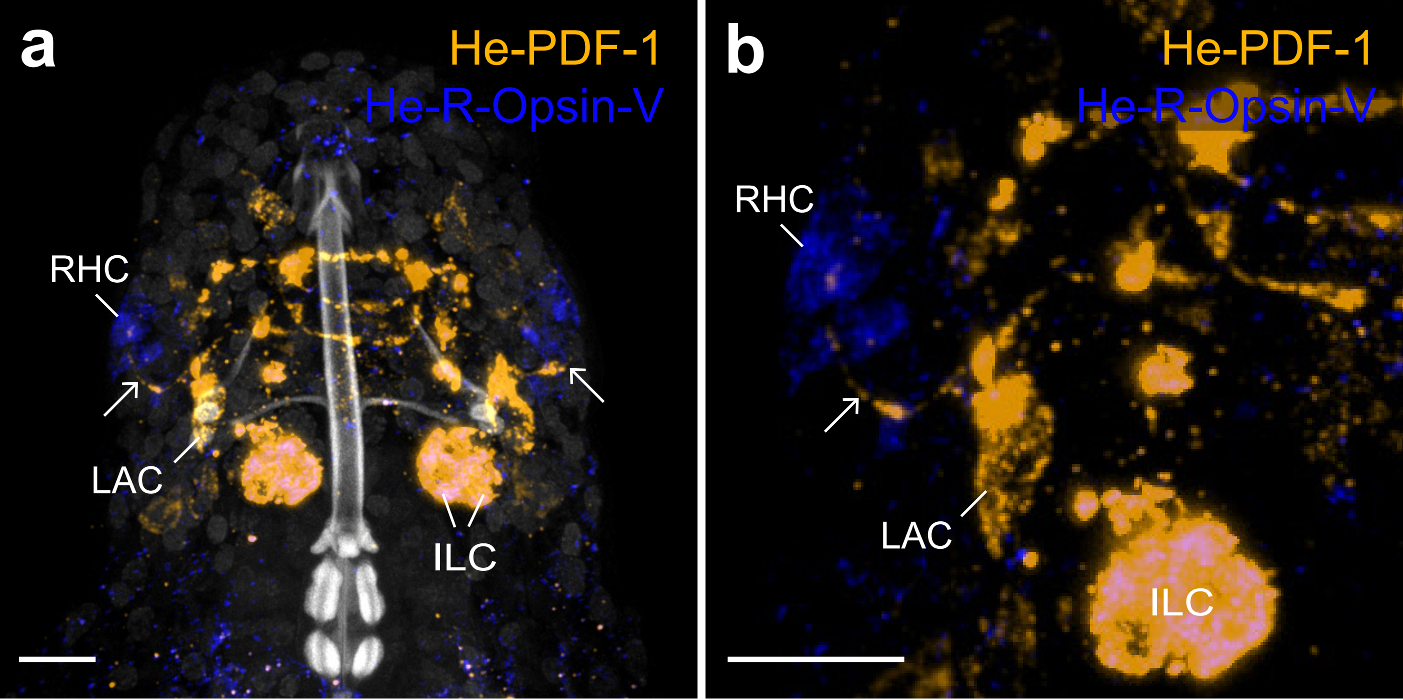


supplementary Fig. 4

### suppl_fig_5.pdf

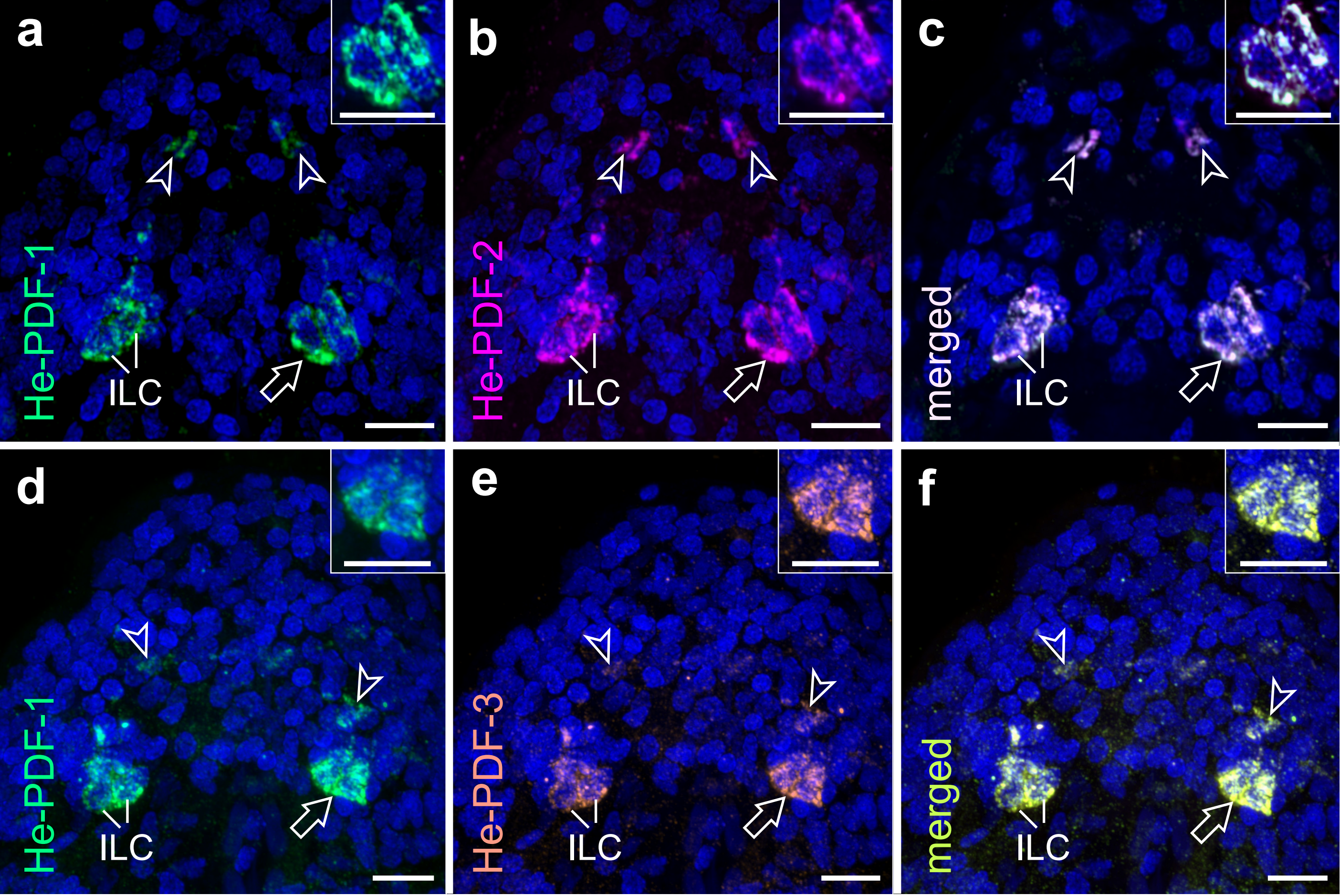


supplementary Fig. 5

### suppl_fig_6.pdf

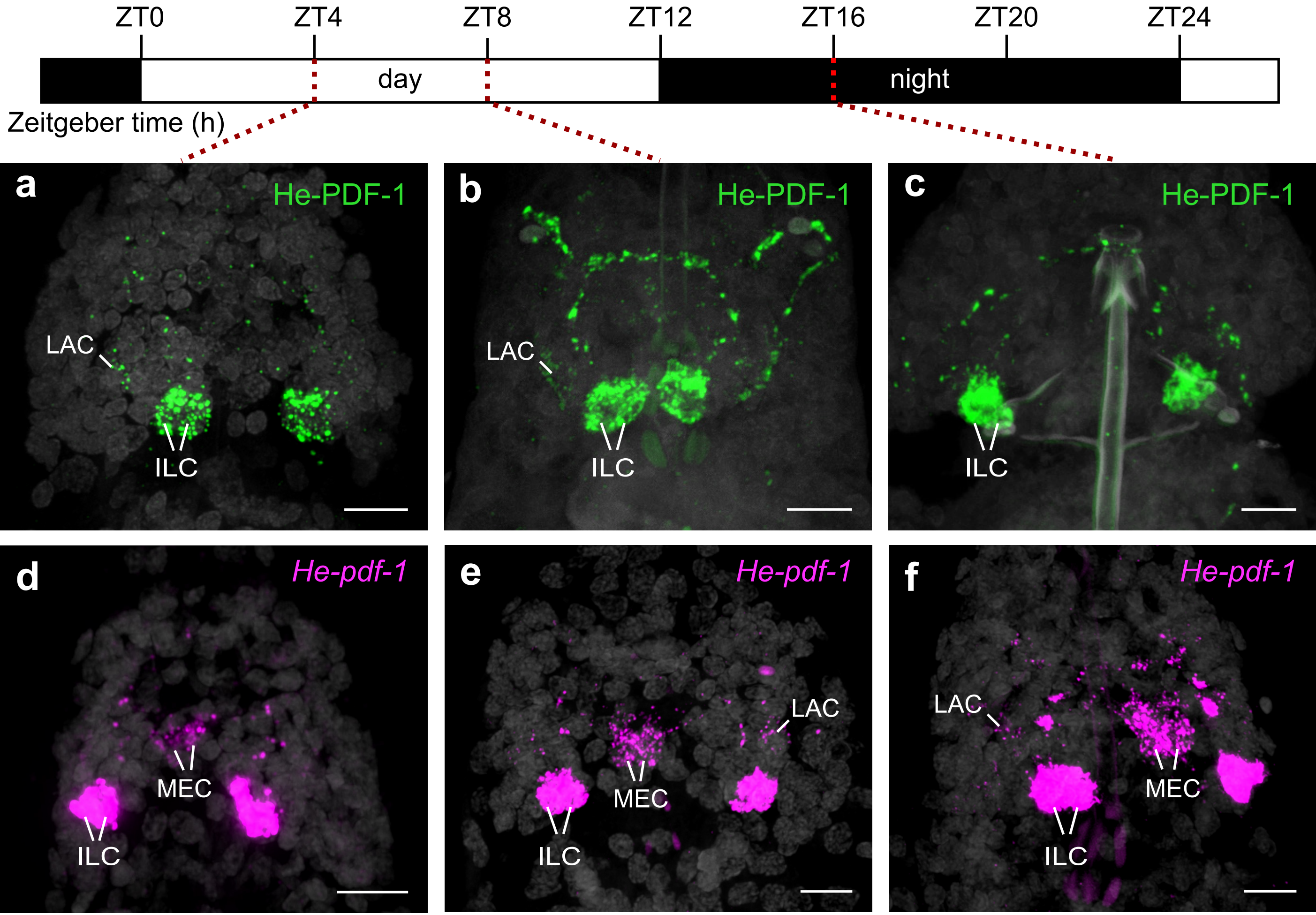


supplementary Fig. 6

### suppl_fig_7.pdf

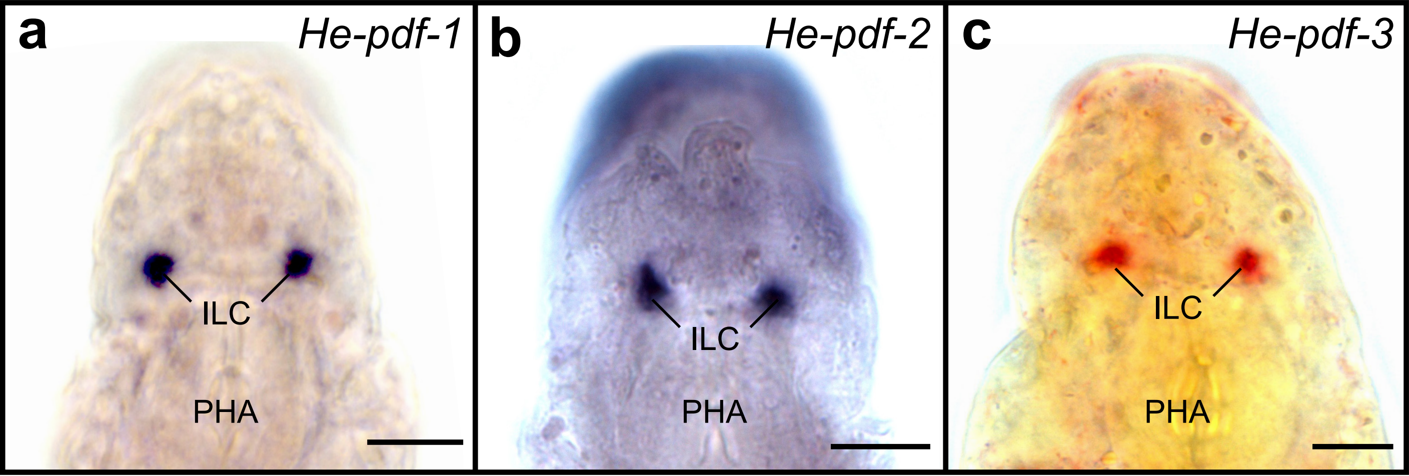

supplementary Fig. 7

### suppl_fig_8.pdf

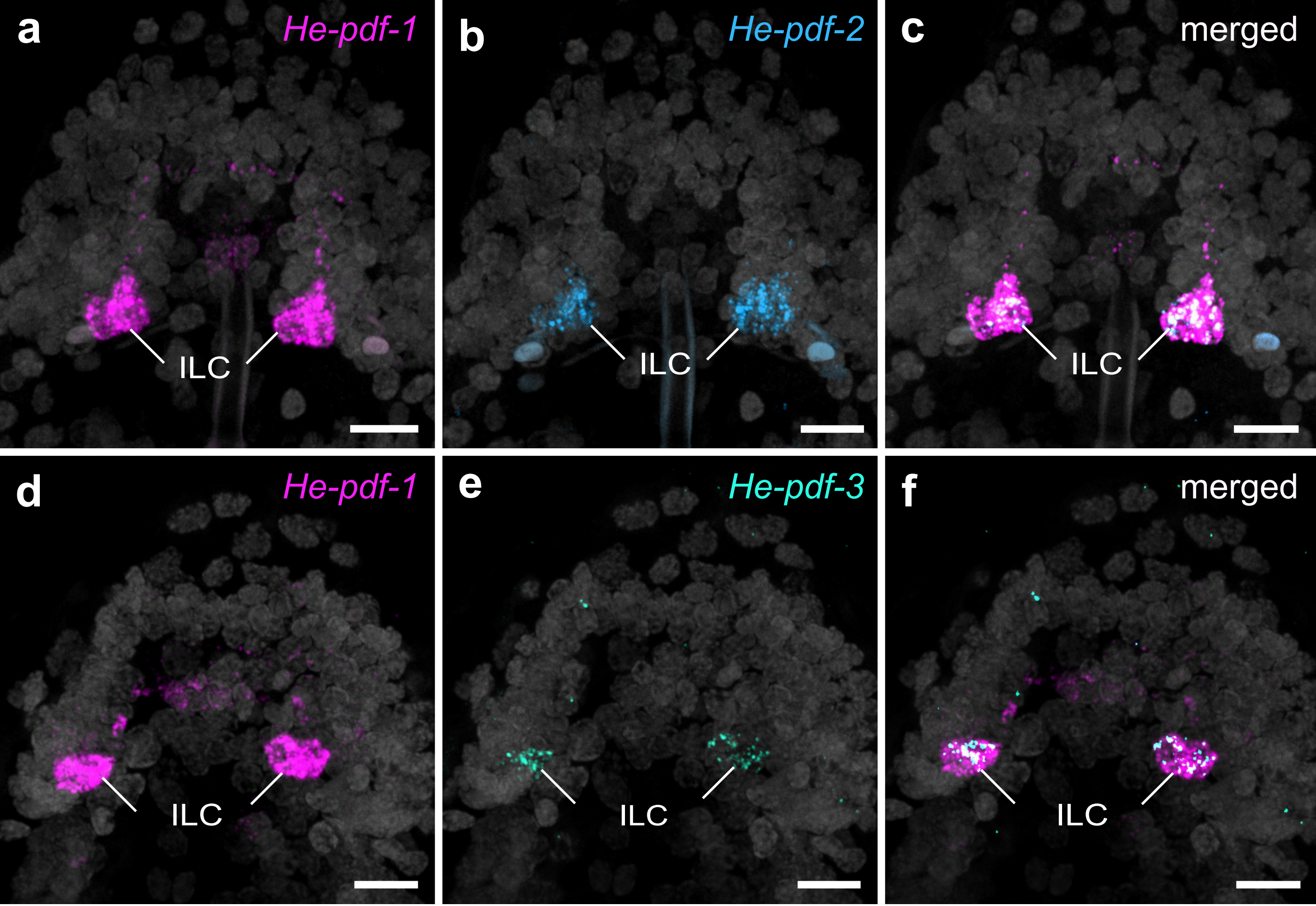


supplementary Fig. 8

### suppl_fig_9.pdf

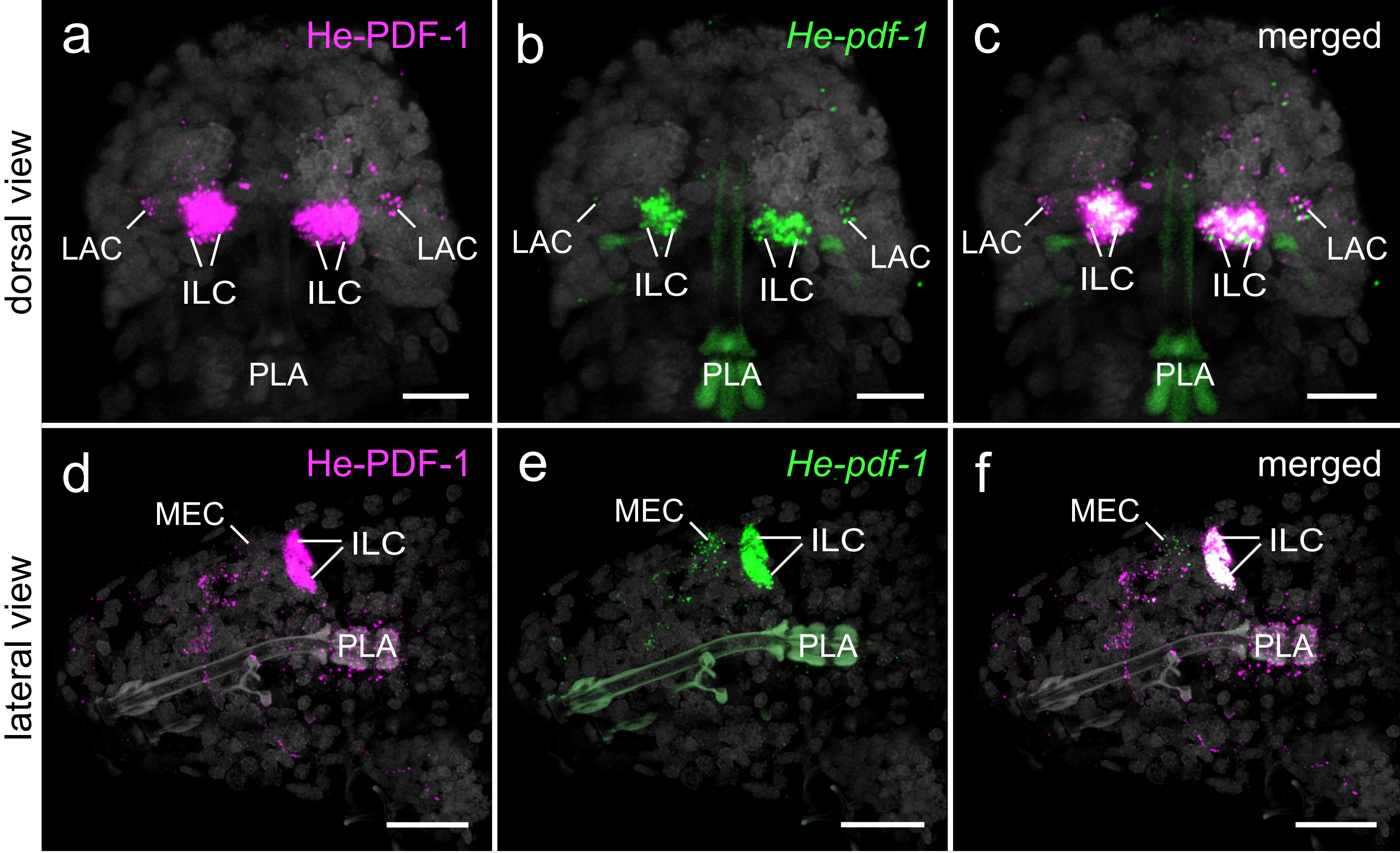


supplementary Fig. 9

### suppl_fig_11.pdf

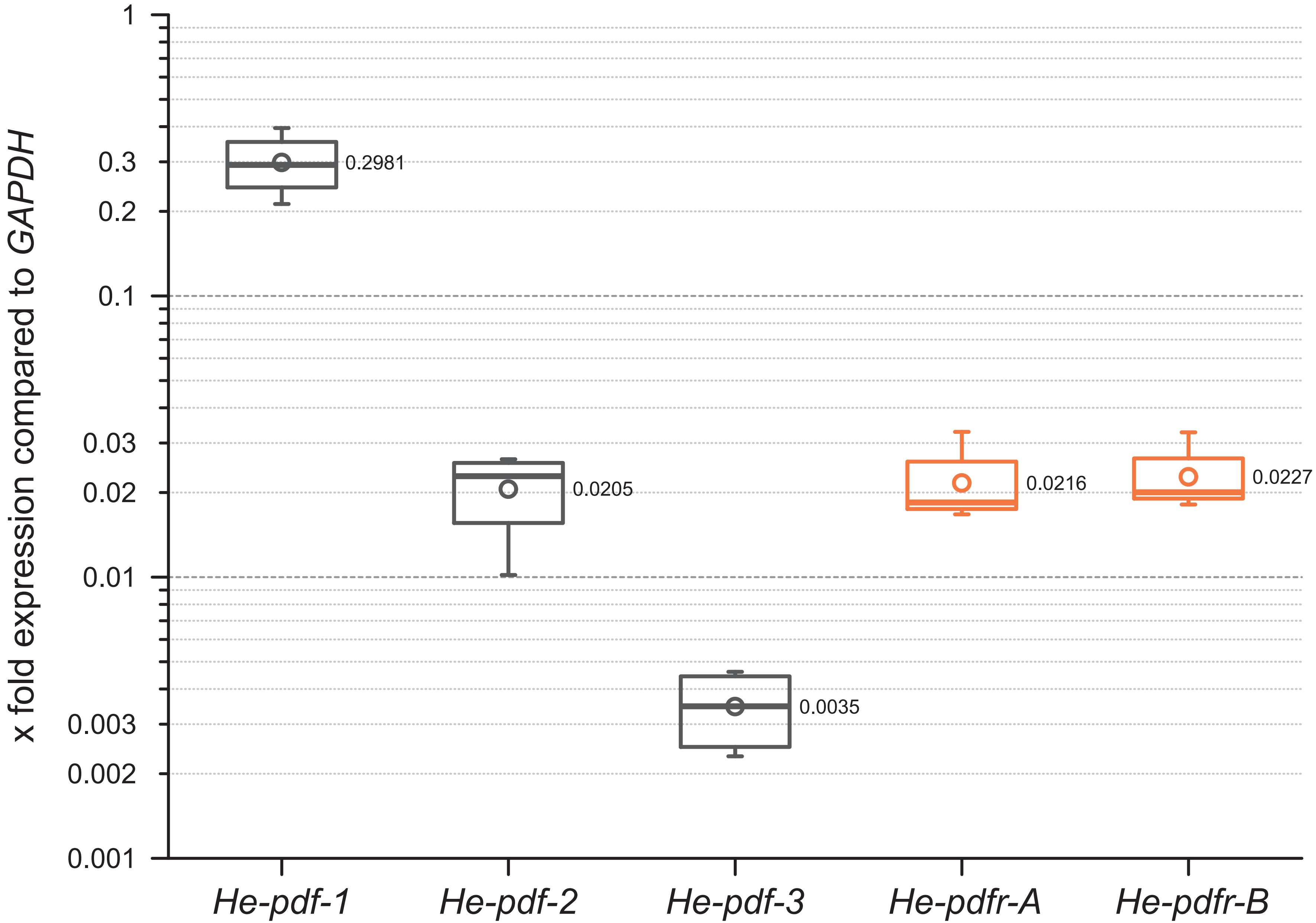


supplementary Fig. 11

### suppl_fig_12.pdf

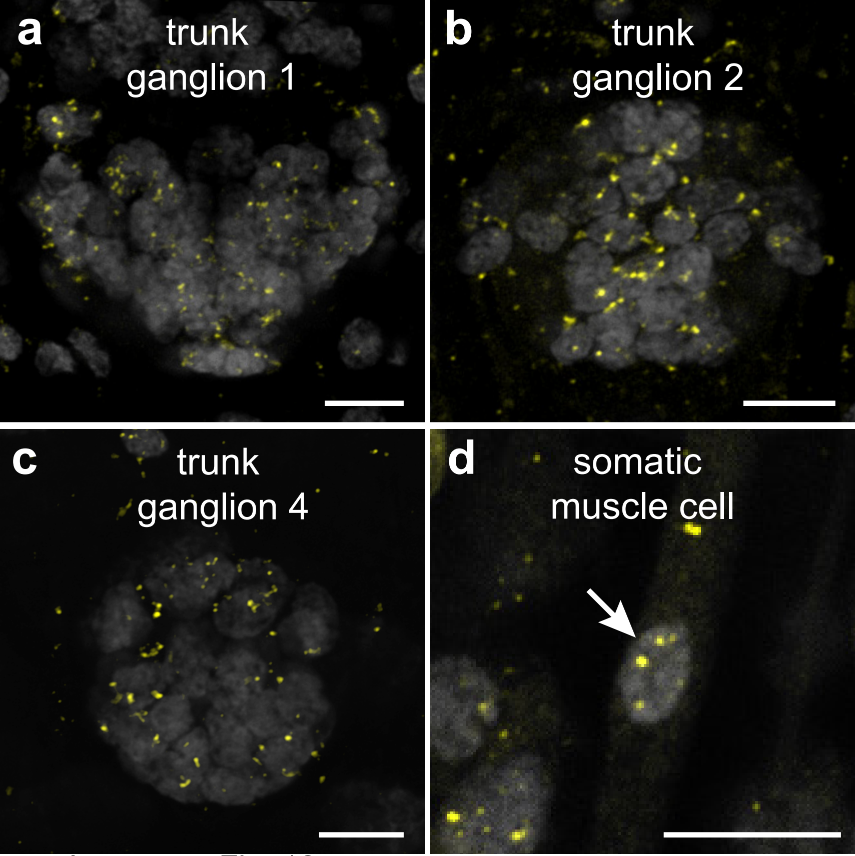


supplementary Fig. 12

### suppl_fig_13.pdf

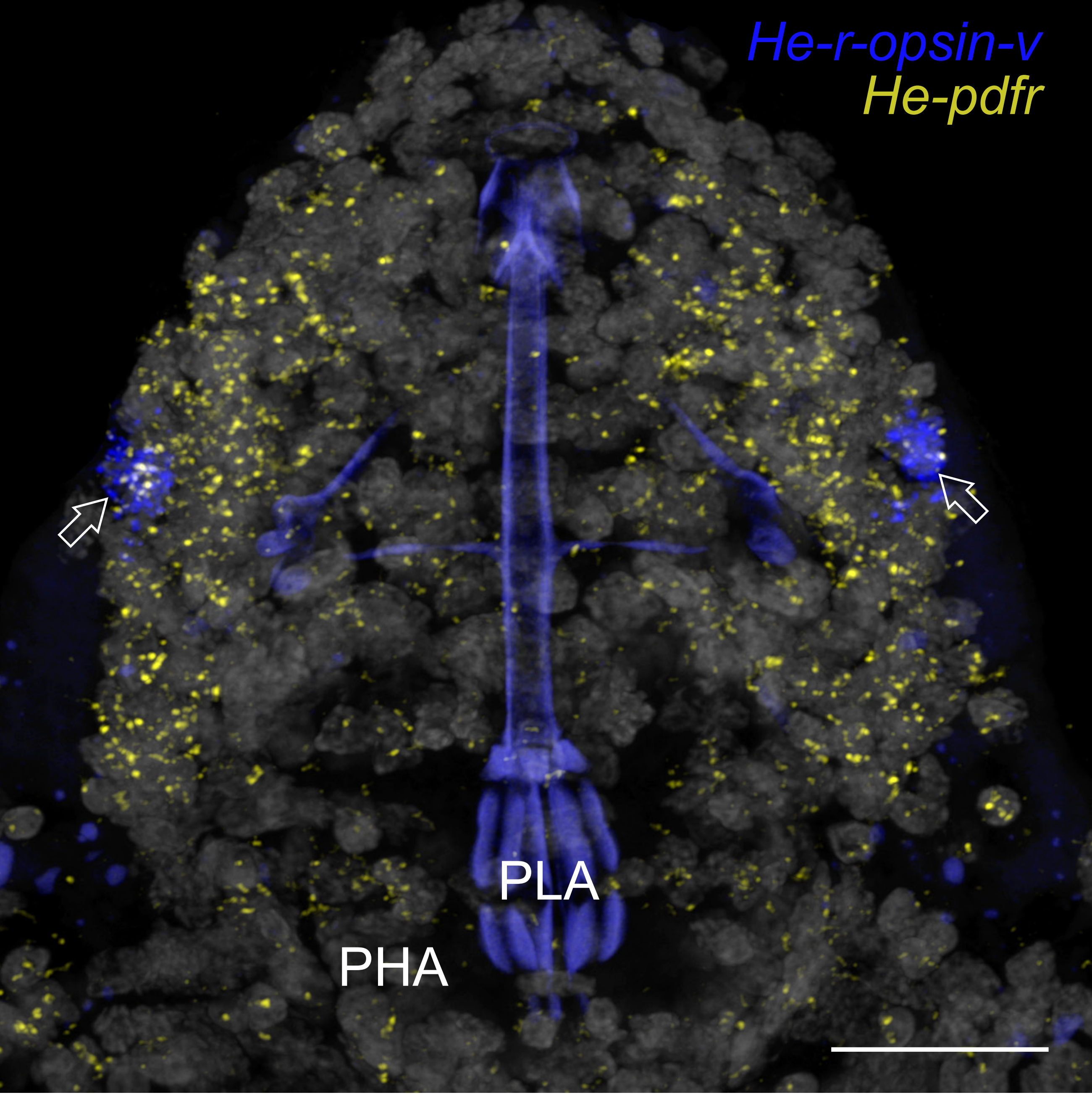


supplementary Fig. 13

### suppl_fig_14.pdf

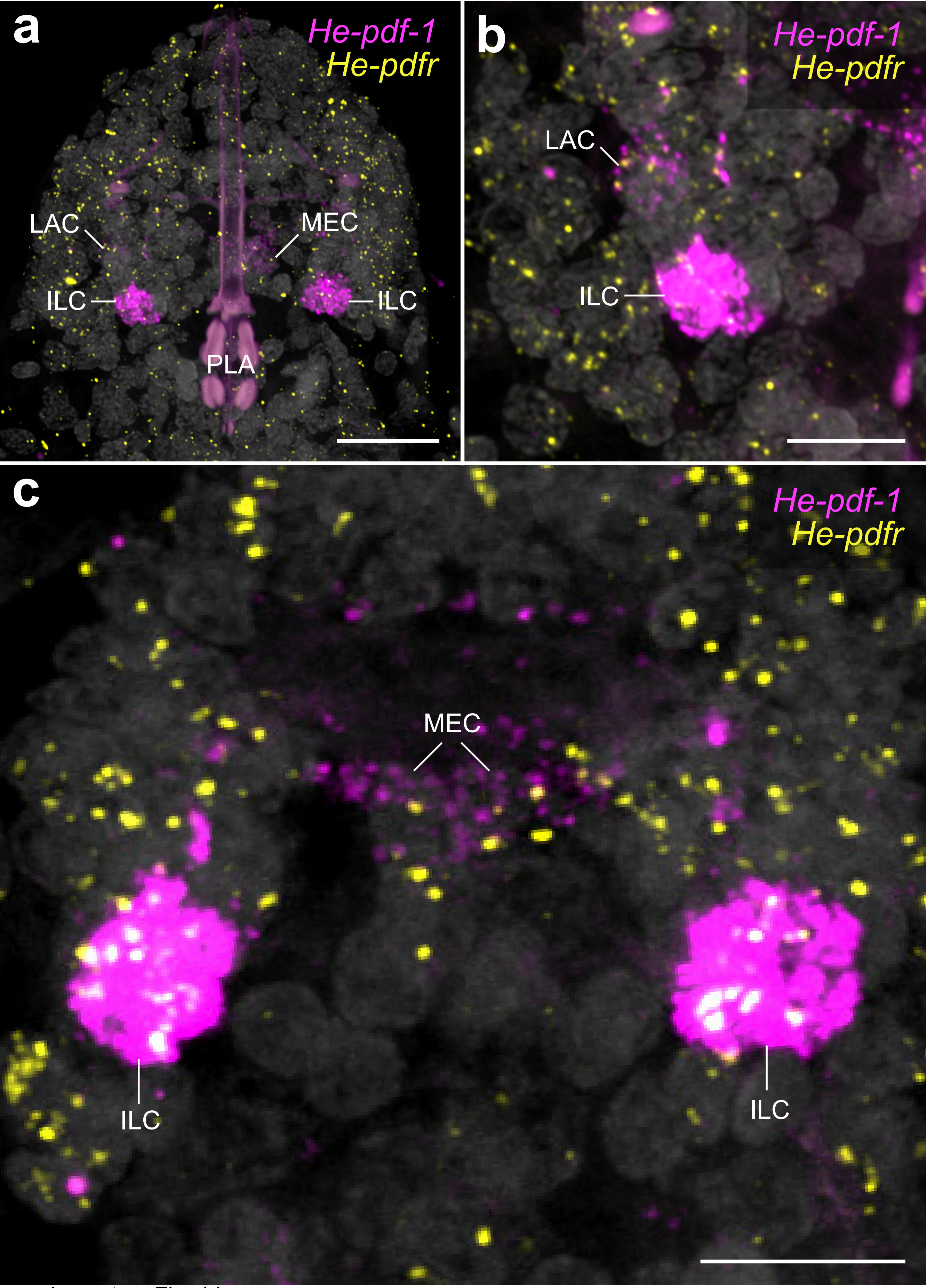


supplementary Fig. 14

### suppl_fig_15.pdf

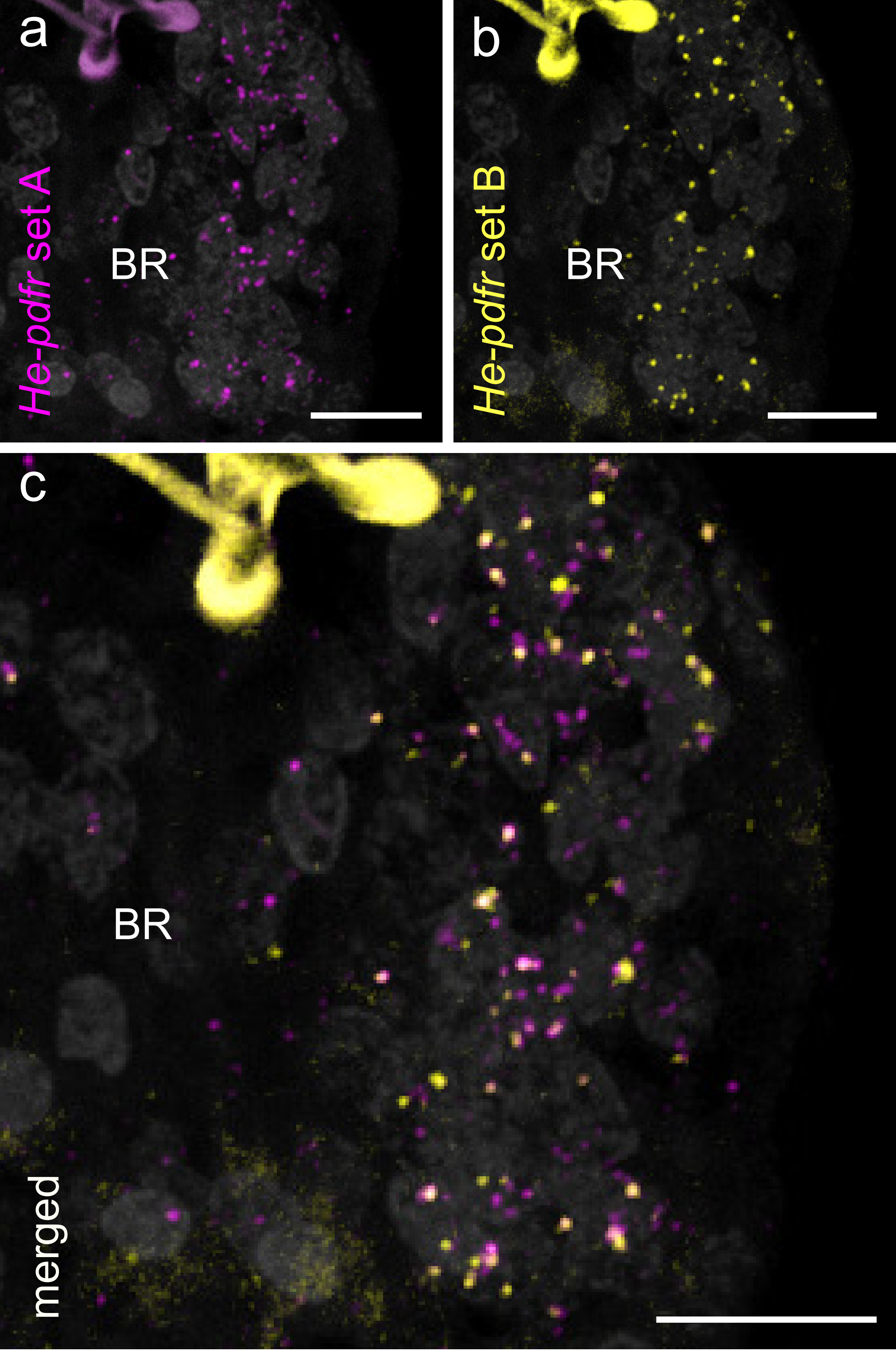


supplementary Fig. 15
